# Faithful Modeling of Terminal CD8 T Cell Dysfunction and Epigenetic Stabilization *In Vitro*

**DOI:** 10.1101/2025.06.22.660847

**Authors:** Amir Yousif, Abbey A. Saadey, Ava Lowin, Asmaa M. Yousif, Ankita Saini, Madeline R. Allison, Kelley Ptak, Eugene M. Oltz, Hazem E. Ghoneim

## Abstract

Epigenetic scarring of terminally dysfunctional CD8 T cells hinders long-term protection and response to immune checkpoint blockade during chronic infections and cancer. We developed a faithful *in vitro* model for CD8 T cell terminal dysfunction as a platform to advance T cell immunotherapy. Using TCR-transgenic CD8 T cells, we found that 1-week peptide stimulation, mimicking conditions in previous models, failed to induce a stable exhaustion program in CD8 T cells. In contrast, prolonged stimulation for 2-3 weeks induced T cell dysfunction but triggered activation-induced cell death, precluding long-term investigation of exhaustion programs. To better mimic *in vivo* exhaustion, we provided post-effector, chronic TGFβ1 signals, enabling survival of chronically stimulated CD8 T cells for over 3 weeks. These conditions induced a stable state of terminal dysfunction (T_Dysf_), marked by a stable loss of effector, cytotoxicity, and memory programs, along with mitochondrial stress and impaired protein translation. Importantly, transcriptomic and epigenetic analyses confirmed the development of terminal exhaustion-specific signatures in T_Dysf_ cells. Adoptive transfer of T_Dysf_ cells revealed their inability to recall effector functions or proliferate after acute LCMV rechallenge. This novel tractable model system enables investigation of molecular pathways driving T cell terminal dysfunction and discovery of new therapeutic targets for cancer or chronic infections.

## Introduction

CD8 T cell effector function is critical for clearing viral infections or spontaneously arising tumor cells. When antigen-primed CD8 T cells experience prolonged antigenic stimulation and inflammatory microenvironments, they progressively lose their capacity to kill virus-infected or tumor cells, becoming dysfunctional. This state of T cell dysfunction, also called exhaustion, is characterized by high expression of surface inhibitory receptors, as well as a gradual loss of effector functions, proliferation capacity, and memory potential, along with impaired mitochondrial function and protein translation (1, 2). The progressive nature of T cell dysfunction coupled with variations in spatial cues they receive produce heterogeneous populations of exhausted T cells (TEX) (2–5). These populations include a progenitor subset that retains self-renewal capacity and effector programs; a cytolytic subset that exhibits the highest cytotoxic function among TEX subsets; and a terminally exhausted subset. The latter develops after persistent stimulation and has the most compromised effector functions, poorest survival and proliferative capacities, and significant mitochondrial dysfunction (1–3, 6). Immune checkpoint blockade (ICB) is a revolutionary cancer immunotherapy that reinvigorates both the progenitor and cytolytic TEX subsets by blocking signals from inhibitory receptors (IRs), such as CTLA-4 and PD-1. However, terminally exhausted T cells remain refractory to ICB—a major obstacle for achieving efficient ICB responses in many patients with cancer.

We and others have shown that the epigenome of TEX cells plays a critical role in locking their fate commitment (3, 7–14). Importantly, de novo DNA methylation enforces the silencing of effector and memory programs in TEX cells, restraining their response to ICB therapy (8). Recent studies have revealed key roles for various microenvironmental signals in regulating T cell dysfunction, such as the strength and duration of TCR stimulation, chronic TGFβ1 exposure, type I IFN signaling, hypoxia, and limited nutrients (1, 2). Additionally, recent studies revealed a critical role for hypoxia—adapted for *in vitro* modeling (15)—in driving and potentially accelerating T cell dysfunction (16, 17). Yet the specific signals critical for initiating and imprinting de novo epigenetic programs that induce terminal exhaustion in CD8 T cells remain unclear. This major gap in the field is due to the diverse range of cues that CD8 T cells encounter *in vivo* within tumor or chronically infected tissue microenvironments. This complexity makes it challenging to isolate and understand the individual and combined effects of exhaustion-inducing signals on epigenetic regulation of TEX cells.

As such, reductionist approaches will be critical for dissecting how signals in each microenvironment drive or counteract epigenetic programming of T cell exhaustion. Recent efforts to develop *in vitro* models of T cell exhaustion have aimed to address these challenges (18–21). In previous models, CD8 T cells acquire exhaustion-like features, including upregulation of inhibitory receptors, reduced proliferation, and mitochondrial stress. Yet, several models primarily utilized persistent TCR stimulation as the sole driver of T cell exhaustion, with repeated antigen-dependent or independent (anti-CD3) stimulation under a short timeline of 5-10 days (18–21). The fundamental obstacle we have found in TCR-based approaches is that repeated TCR stimulations of CD8 T cells leads to activation-induced cell death (9, 20). Thus, persistent TCR signaling alone hinders the ability to replicate the developmental timeline of T cell exhaustion observed *in vivo*, in which at least 2-3 weeks of chronic stimulation in the presence of other microenvironmental signals are necessary to establish a terminally dysfunctional state during chronic infection or cancer (11, 22–24). Furthermore, it remains critical to track whether the dysfunctional state established *in vitro* is stable after antigen stimulation is withdrawn—a core defining feature of the epigenetically fixed state of terminal exhaustion observed in chronic virus- or tumor-specific CD8 T cells *in vivo* (11, 22–24). Therefore, it is essential to develop a robust model system that faithfully recapitulates the progressive nature of T cell exhaustion and enables tracking of chronically stimulated T cells over extended periods. Such a system is vital for developing reliable discovery platforms to identify novel therapeutic targets aimed at preventing or reversing T cell dysfunction and enhancing the effectiveness of T cell immunotherapies.

Here, we developed a new antigen-dependent model system of T cell exhaustion using TCR-transgenic, GP33-specific CD8 T cells. We found that prolonged (2-3 weeks) rather than short-term antigen stimulation (1 week) was required to induce stable CD8 T cell exhaustion. However, as with other systems, activation-induced cell death limited our long-term studies of chronically stimulated CD8 T cells. To better mimic *in vivo* conditions, chronic TGFβ1 signals were introduced at the post-effector phase, which supported antigen-specific CD8 T cell survival for 3 weeks during chronic stimulation. The combined signals and long-term viability faithfully induced terminal dysfunction in CD8 T cells (T_Dysf_), which displayed a remarkable loss of effector and memory programs, mitochondrial stress, and impaired protein translation. Transcriptome and epigenome analyses of these dysfunctional lymphocytes confirmed the acquisition of exhaustion-specific signatures. Furthermore, adoptive T cell transfer studies showed that T_Dysf_ cells had impaired effector recall responses and proliferation capacity. This novel model enables investigation of the long-term developmental process that leads to terminal dysfunction in antigen-specific CD8 T cells, providing a faithful platform for identifying new therapeutic targets to block or reverse epigenetic processes stabilizing terminal dysfunction.

## Results

### Acute antigenic stimulation fails to induce a dysfunctional program in CD8 T cells

To determine whether repeated TCR stimulation for one week can induce a stable dysfunctional program in CD8 T cells, we utilized naïve TCR-transgenic murine CD8 T cells “P14 cells” that can be stimulated by LCMV-derived GP33 peptide. We compared P14 cells that underwent an initial activation using anti-CD3 plus anti-CD28 stimulation on day 0-2 (“Acute-2d”) to those that received repeated GP33 peptide stimulations following this activation phase until day 7 (“Acute-7d”) (**Figure 1A**). We first characterized the phenotype of P14 CD8 T cells following the initial activation phase (days 0-2) and observed that over 98% of P14 cells upregulated CD44 and PD-1, with the majority co-expressing PD-1 and Tim-3—indicating robust activation and effector differentiation (**Fig. 1B-D, Supplemental Fig.1B**). By day 7, repeated GP33 stimulation led to sustained high frequencies of CD44+ PD-1+ and PD-1+ Tim-3+ populations in Acute-7d P14 cells compared to 2d-stimulated cells (**Figure 1C-D, Supplemental Fig. B**). This pattern mirrors the *in vivo* upregulation of surface inhibitory receptors on activated effector CD8+ T cells (25). To track whether these observed features are stable after stopping antigen stimulation, we maintained antigen-experienced P14 cells for 12 additional days without GP33 stimulation, then rechallenged them with GP33 peptide on days 12 and 19 (i.e., 5 and 12 days after GP33 antigen removal) (**Figure 1A**). The frequencies of CD44+ PD-1+ and PD-1+ Tim3+ populations within Acute-7d P14 cells gradually declined over time but remained elevated compared to Acute-2d cells by day 19 (**Figure 1C-D**, **Supplemental Figure 1B-D**). These findings suggest that Acute-7d stimulated P14 cells do not have stable maintenance of IR expression following rest from antigen stimulation, contrasting them from exhausted T cells *in vivo*, which retain high levels of IRs (26, 27).

**Figure 1.**
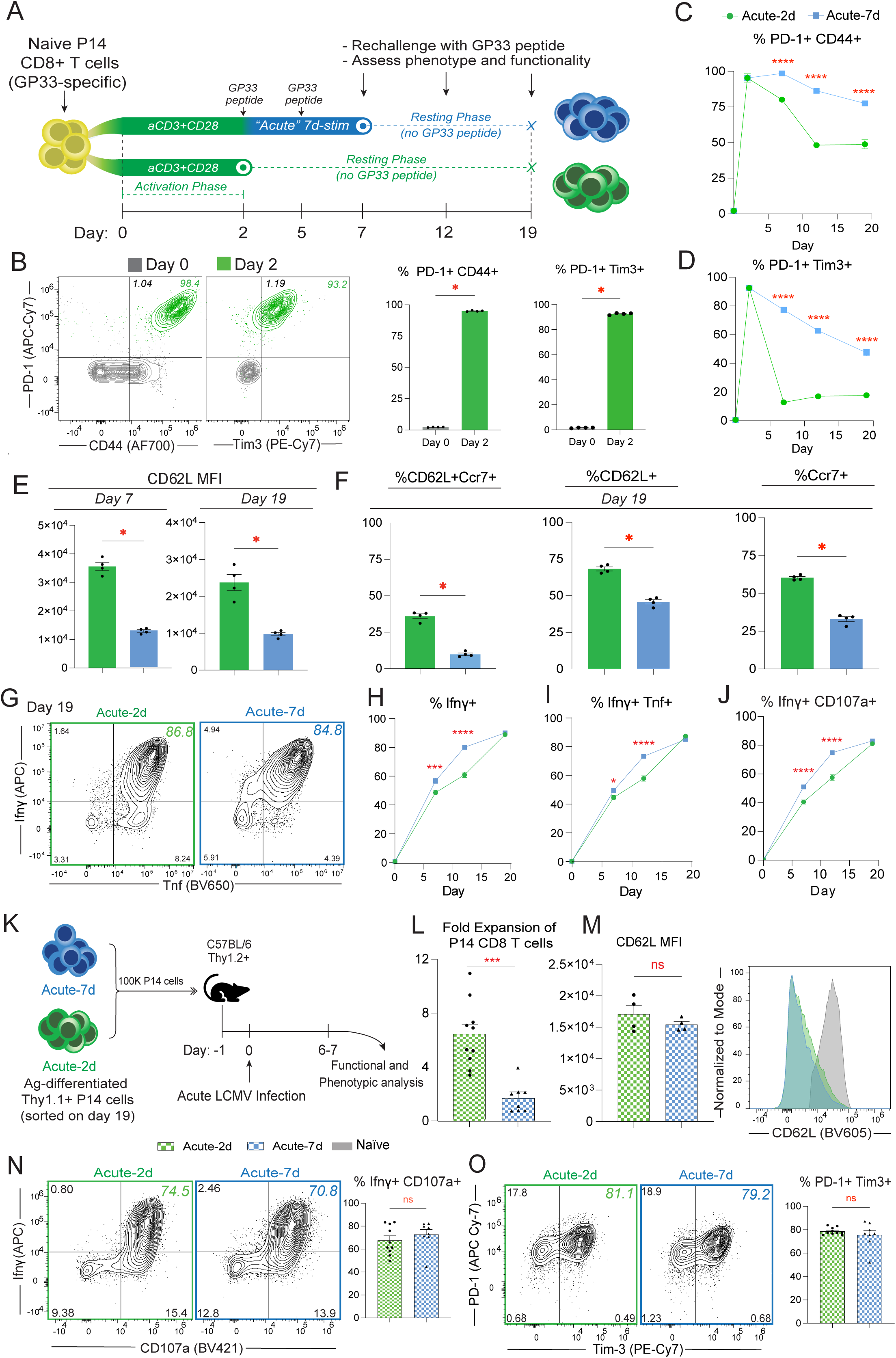
Acute antigenic stimulation for one week fails to induce a dysfunctional program in CD8 T cells. (**A**) Schematic for *in vitro* stimulation of GP33-specific P14 CD8+ T cells after isolation from spleens of naïve P14 mice. P14 cells were first activated with anti-CD3 and anti-CD28 from day 0-2, followed either by resting until day 19 (“Acute-2d” condition), or by two rounds of GP33 peptide stimulation from day 2-7 followed by resting until day 19 (“Acute-7d” condition). (**B**) Representative FACs plots confirming the activation of P14 cells from day 0 to day 2 shown by PD-1, CD44, and Tim3 expression. Summery bar graphs depict the frequency of PD-1+ CD44+ as well as PD-1+ Tim3+ P14 cells in naive (Day 0) and stimulated P14 cells (Day 2). (**C**) Longitudinal tracking of %PD-1+ CD44+ P14 cells for either Acute-2d (bottom line, green) or Acute-7d (top line, blue). (**D**) Longitudinal tracking of %PD-1+ Tim3+ P14 cells for either Acute-2d (bottom line, green) or Acute-7d (top line, blue). (**E**) Expression levels (as shown by mean fluorescence intensity) of CD62L from in vitro cultured Acute-2d and Acute-7d P14 cell on days 7 and 19 of stimulation. (**F**) Frequencies of CD62L+ Ccr7+; total CD62L+; and total Ccr7+ in vitro cultured Acute-2d and Acute-7d P14 cells on Day 19. **(G)** Representative FACS plots of Ifnγ and Tnf expression on P14 cells on day 19 after GP33 peptide rechallenge. **(H)** Longitudinal tracking of % total Ifnγ+; **(I) %** Ifnγ+ Tnf+; and **(J) %** Ifnγ+ CD107a+ P14 cells after GP33 peptide rechallenge at the indicated timepoints. **(K)** Schematic for adoptive transfer of 100K “Acute-2d” (green) or “Acute-7d” (blue) Thy1.1+ P14 cells generated *in vitro* into C57BL/6 mice on day -1, followed by acute LCMV infection on day 0, and then functional and phenotypic analyses of adoptively transferred P14 cells on day 6-7 p.i. **(L)** Bar graph depicting the fold expansion of P14 CD8+ T cells relative to the endogenous GP33+ CD8 T cells for adoptively transferred acute 2-day and acute 7-day P14 cells. **(M)** Expression levels (MFI) of CD62L in adoptively transferred 2-day and acute 7-day P14 cells along with a representative histogram. **(N)** Representative FACS plots and summary bar graph of % Ifnγ+ CD107a+ P14 cells after *ex-vivo* GP33 peptide restimulation of splenocytes. **(O)** Representative FACS plots and summary bar graph of % PD-1+ Tim-3+ P14 cells on day 6 p.i. For **B-J** and **M** All *n*□=□4 biological replicates, representative of two to three independent experiments. For **L, N** and **O**, data were pooled from two independent experiments with *n*□=□3–4 biological replicates per group for each experiment. Adjusted P value \**P<0.05*, \*\**P*□*<*□*0.01*, \*\*\**P*□*<*□*0.001*, \*\*\*\**P*□*<*□*0.0001*. Comparisons were determined by Two-way ANOVA (**C, D, H-J**), or the Mann–Whitney U test (unpaired, two sided) (**B, E, F, L-O**). Error barsLindicate meanL±LSEM.

A key characteristic of T cell exhaustion is the failure to restore memory programs after antigen stimulation is removed (11–14, 24). To determine whether repeated antigen stimulation for one week impacts the recovery of memory programs after antigen withdrawal, we tracked expression levels of memory and stemness-associated programs before and after resting, specifically monitoring Ly108, L-selectin (CD62L), Ccr7, and IL-7 receptor (CD127). Importantly, Acute-7d P14 cells initially downregulated Ly108 and CD62L compared to Acute-2d cells on day 7; however, they recovered similar expression levels of Ly108 after resting, while Acute-2d cells maintained higher CD62L expression (**Figure 1E**, **Supplemental Figure 1I**). We also observed that the frequencies of CD62L+ Ccr7+ cells remain high in Acute-2d cells as compared to Acute-7d cells after resting, while both Acute-2d and Acute-7d cells maintained high expression of Il7r on day 19 (**Figure 1F, Supplemental Figure 1J**). Notably, there were also no significant differences in the proliferative activity (measured by Ki67 expression) between the two groups on day 19 (**Supplemental Figure 1H**). These findings suggest that one week of repeated antigen stimulation does not lead to a stable loss of memory features, mirroring the retained memory potential in effector CD8+ T cells isolated after one week of chronic LCMV infection (24).

Next, we tracked changes in effector function of Acute-2d and Acute-7d P14 cells using GP33 antigen rechallenge at different timepoints. Here, we found both Acute-2d and Acute-7d cells exhibited profound polyfunctionality as indicated by Ifnγ and Tnf co-expression, as well as high degranulation activity (CD107a) (**Figure 1G-J, Supplemental Figure 1G**). Notably, the frequencies of polyfunctional cells continued to increase during the resting period (**Figure 1H-J**). We also found that Acute-7d P14 cells had significantly higher degranulation activity and increased expression of cytotoxic molecules Gzmb and Perforin compared to Acute-2d (**Supplemental Figure 1E-G**). These observations indicate that repeated acute antigen stimulation may be necessary to fully establish cytolytic effector differentiation in CD8 T cells. Additionally, we observed relatively less cellular apoptosis in Acute-2d compared to Acute-7d P14 cells (Annexin V staining), while both maintained substantial viable populations after resting (**Supplemental Figure 1K**). Finally, there was no significant difference in the mitochondrial mass (MitoTracker-Green) or oxidative phosphorylation activity as probed by measuring mitochondrial reactive oxygen species “ROS” levels (MitoSox) (**Supplemental Figure 1L-M**). Overall, our findings demonstrate that one week of *in vitro* antigen stimulation alone is insufficient to induce stable dysfunction, suggesting that longer duration of antigenic stimulation and/or additional microenvironmental signals are necessary to establish a terminally exhausted-like profile in CD8 T cells.

Lastly, to confirm whether one week of *in vitro* antigen stimulation is insufficient to induce features of exhaustion, we assessed the recall responses of resting Acute-2d or Acute-7d CD8 T cells by adoptively transferring them into mice followed by acute LCMV infection, as terminally exhausted T cells have limited capacity to mount a response to antigen rechallenge (11, 24). Here, we sorted congenically distinct Thy1.1+ P14 cells on day 19 of *in vitro* culture, then adoptively transferred 100-150K cells into C57BL/6 (Thy1.2+) animals. One day later, we challenged these chimeric animals with acute LCMV infection and assessed recall responses of P14 cells after 6-7 days of infection (**Figure 1K**). Both Acute-2d and Acute-7d P14 cells underwent extensive antigen-dependent proliferation; however, Acute-2d cells exhibited significantly greater expansion than Acute-7d cells (**Figure 1L**), potentially due to enhanced survival and/or L-selectin-mediated homing to lymphoid tissues of the adoptively transferred Acute-2d cells (**Figure 1E, Supplemental Figure 1K**). Yet, both Acute-2d and Acute-7d cells showed comparable CD62L expression following infection-induced expansion **(Figure 1M**). Next, we assessed the functionality of these *in vivo*-expanded P14 cells via *ex vivo* GP33 peptide stimulation. We found that both Acute-2d and Acute-7d P14 cells exhibited heightened polyfunctionality, indicated by elevated expression levels of effector molecules such as Ifnγ, CD107a, Perforin, Gzmb, and Tnf (**Figure 1N, Supplemental Figure 2C-D, G**). We also observed high levels of memory, stemness, proliferation, and activation markers after rechallenge, including CD62L, Ki67, CD44, Ly108, Tcf1, PD-1, Tim3 (**Figure 1O, Supplemental Figure 2A-B, E-I)**. Of note, similar expression levels of these markers were also observed within the peripheral organs, including liver and lungs (**Supplemental Figure 2J-S**). Overall, these findings confirm that repeated antigen stimulation of CD8 T cells for one week fails to induce a terminal dysfunctional program, as these cells retain heightened recall responses during acute infection.

### Chronic antigenic stimulation faithfully induces terminal dysfunction

To develop a model system for terminal dysfunction in CD8 T cells, we extended the duration of repeated GP33 peptide stimulation for an additional two weeks (“Chronic GP33 stim”) (**Figure 2A**). Compared to acutely stimulated “Acute-7d” condition, 2-3 weeks of prolonged antigen stimulation produced a higher frequency of PD-1+ Tim3+ P14 cells (**Figure 2B**). However, the chronically stimulated P14 cells exhibited substantial loss of viability, with a progressive decline in total cell numbers and increased cell death, likely due to activation-induced cell death—a major limitation for studying CD8 T cells under prolonged periods of TCR stimulation *in vitro* (20) (**Figure 2C, Supplemental Figure 3F**).

**Figure 2.**
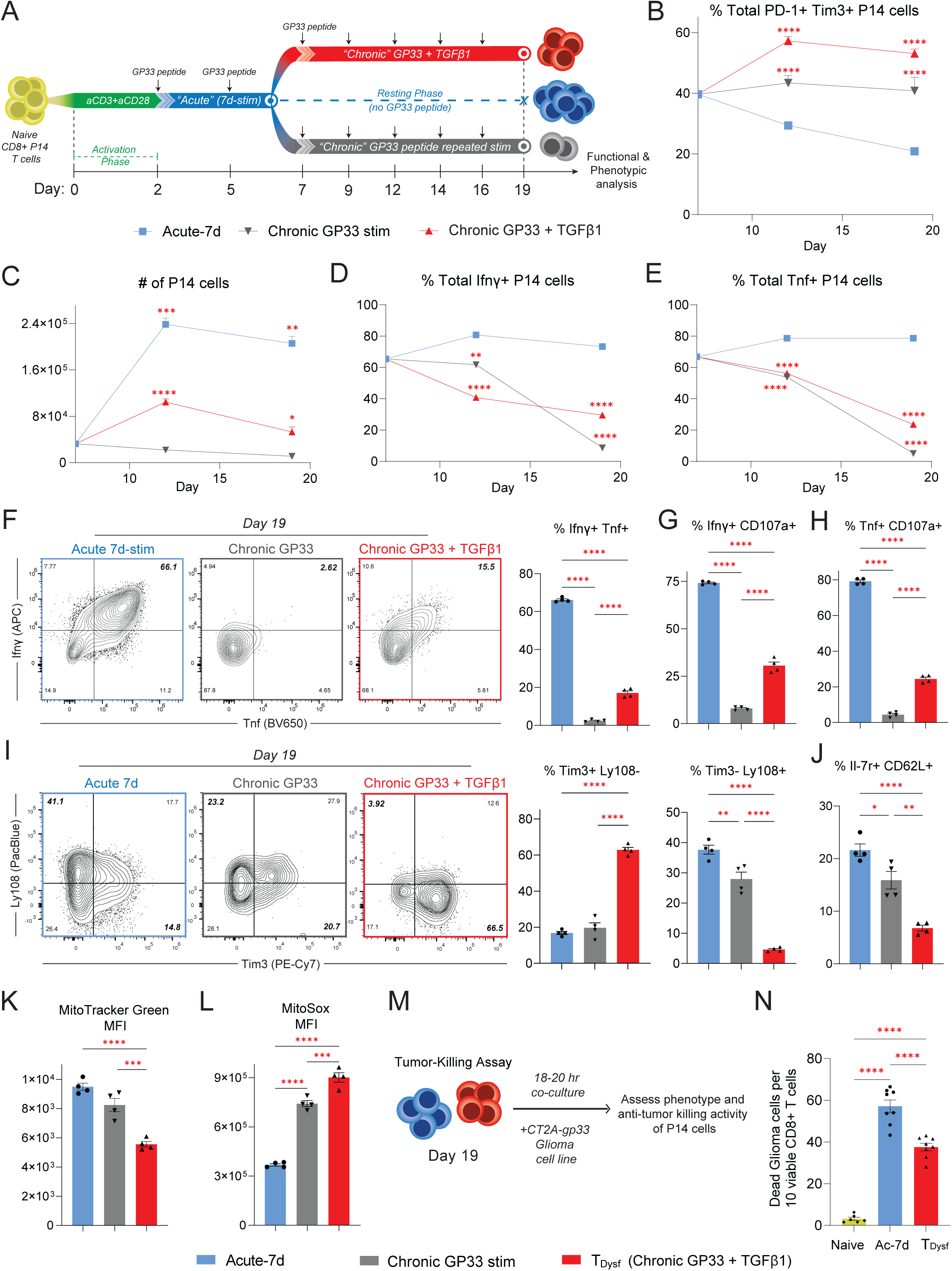
Chronic antigenic stimulation faithfully induces terminal dysfunction. (**A**) Schematic for *in vitro* stimulation of P14 cells—activated P14 cells were acutely stimulated by GP33 peptide from day 2-7 and resting until day 19 (“Acute-7d” condition—blue), or repeatedly GP33-stimulated from day 2-19 with chronic TGFβ1 exposure (day 7-19—red “T_Dysf_”) or without TGFβ1 signals (gray). Longitudinal tracking of (**B**) frequencies of PD-1+ Tim3+ P14 cells, (**C**) P14 cell numbers/200 µl, (**D**) %Ifnγ+ and (**E**) %Tnf+ from day 7-19 for Acute-7d (blue), Chronic GP33 stim (gray) or Chronic GP33+TGFβ1 (red). (**F**) Representative FACS plots and bar graphs of Ifnγ and Tnf expression, (**G**) %Ifnγ+ CD107a+ or (**H**) %Tnf+ CD107a+ P14 cells on day 19 after GP33 peptide rechallenge. (**I**) Representative FACS plots and bar graphs of Tim3 and Ly108 expression. (**J**) Summary bar graph of % Il-7r+ CD62L+ P14 cells on day 19. Expression level (gMFI) of (**K**) MitoTracker Green dye (for mitochondrial mass) and (**L**) MitoSox Red dye (for mitochondrial ROS) within P14 cells on day 19. (**M**) Schematic for tumor-killing assay of naive “yellow”, Acute-7d, or T_Dysf_ P14 cells co-cultured with GP33-expressing CT2A glioma tumor cells. (**N**) Bar graph showing numbers of dead CT2A-GP33 cells per 10 viable P14 cells. All *n*□=□4 biological replicates, representative of two to three independent experiments. Adjusted P value \**P<0.05*, \*\**P*□*<*□*0.01*, \*\*\**P*□*<*□*0.001*, \*\*\*\**P*□*<*□*0.0001*. Comparisons were determined by Two-way ANOVA (**B-E**), or One-way ANOVA (**F-L, N**). Error barsLindicate meanL±LSEM.

Previous studies have attempted to model T cell exhaustion *in vitro* using plate-bound anti-CD3/CD28 stimulation (day 0-2) followed by repeated plate-bound anti-CD3 stimulations for 5-10 days (20, 21). While extended strong TCR stimulation induced high levels of PD-1 and Tim3 (**Supplemental Figure 3A-C**), this method also induced extensive cell death that limits reliable functional assessment and tracking of the epigenetic remodeling towards terminal T cell exhaustion (**Supplemental Figure 3A, D-E**). To better mimic *in vivo* conditions where CD8 T cells can survive and develop exhaustion following prolonged antigen exposure, we provided continuous TGFβ1 signals between days 7 and 19 (“Chronic GP33 stim + TGFβ1”) to ameliorate activation-induced cell death (**Figure 2A**) (9). Our previous work demonstrated that prolonged, post-effector TGFβ1 exposure drives terminal dysfunction in human CD8 T cells that are chronically stimulated in an antigen-independent manner (9). Indeed, we found that chronic GP33 plus TGFβ1-stimulated P14 cells not only showed improved cell viability compared to chronic GP33 alone or plate-bound stimulation approaches (**Figure 2C, Supplemental Figure 3D-F**), but also further increased PD-1 and Tim3 co-expression (**Figure 2B)**. To evaluate the impact of TGFβ1 exposure on CD8 T cell survival, we performed Annexin V and live/dead staining on chronically stimulated CD8 T cells. TGFβ1 supplementation significantly improved the viability of P14 T cells under chronic stimulation, with reduced apoptosis and increased proportions of live cells compared to chronic GP33 stimulation alone on both days 14 and 20 (**Supplemental Figure 3F-H**). These findings suggest that TGFβ1 attenuates activation-induced cell death. The improved viability of P14 cells under chronic GP33 plus TGFβ1 stimulation (hereafter referred to as “T_Dysf_”) enabled robust analysis of exhaustion programming during prolonged antigenic stimulation.

We next examined whether chronic GP33 plus TGFβ1-stimulation induces dysfunction characteristic of terminally exhausted CD8 T cells. By tracking effector cytokine production in chronically *versus* acutely stimulated P14 cells, we observed that chronic GP33 stimulation—with or without TGFβ1—led to a progressive decline in effector cytokine output (**Figure 2D-E**), indicative of a transition toward dysfunction. Importantly, T_Dysf_ exhibited a significant loss of polyfunctionality compared to acutely stimulated cells, with reduced frequencies of Ifnγ+Tnf+, Ifnγ+CD107a+, and Tnf+CD107a+ cells, reflecting severe impairment in both cytokine production and degranulation capacity (**Figure 2F-H**). Furthermore, T_Dysf_ cells showed substantial loss of memory-associated markers, including reduced expression and frequencies of Il7r and CD62L compared to both acutely and chronically stimulated controls (**Figure 2J, Supplemental Figure 4E**), further indicating impaired memory differentiation (**Supplemental Figure 4F**).

We also examined exhaustion-associated surface markers and found that T_Dysf_ cells exhibited a shift toward a terminally exhausted phenotype. Alongside sustained high co-expression of PD-1 and Tim-3, T_Dysf_ cells showed decreased frequencies and expression of Ly108—a marker of the progenitor subset of TEX cells—leading to an increased proportion of Tim3+ Ly108-cells, a phenotype of terminal exhaustion (2, 28, 29) (**Figure 2I**). Furthermore, T_Dysf_ cells demonstrated increased frequencies of Tim3+ CD62L-cells, further supporting their progression toward a terminally exhausted state (**Supplemental Figure 4A**). Notably, T_Dysf_ cells showed elevated expression of CD103, a well-established downstream target gene of TGFβ1 signaling that marks a terminally exhausted subset of human tumor-reactive CD8 T cells (30–33) (**Supplemental Figure 4D**). In addition, T_Dysf_ cells showed increased expression of Gzmb—commonly associated with terminal dysfunction (28, 34)—as well as higher levels of perforin compared to Acute-7d cells, likely reflecting the effects of sustained TCR stimulation (**Supplemental Figure 4B-C**). To assess mitochondrial stress, a key feature linked to terminal dysfunction in CD8+ T cell exhaustion (15, 21, 35), we tracked changes in mitochondrial functions within T_Dysf_ P14 cells and found a significant reduction in mitochondrial mass accompanied by increased mitochondrial ROS levels—a marker of progressive T cell exhaustion *in vivo* (**Figure 2K-L**). These results indicate that chronic antigen plus TGFβ1 stimulation over three weeks establishes a dysfunctional program in CD8+ T cells.

To assess the cytotoxic function of *in vitro*-differentiated P14 cells, we utilized an antigen-dependent tumor-killing assay, in which we co-cultured naïve, chronically or acutely stimulated P14 cells with GP33-expressing CT2A glioma cells for 18-20 hours (**Figure 2M**). Consistent with their impaired effector functions, T_Dysf_ cells demonstrated significantly reduced tumor-killing activity compared to acutely stimulated P14 cells (**Figure 2N**), further demonstrating their exhausted state. Overall, these findings reveal a fundamental role for chronic TGFβ1 signals in driving both human and mouse T cell dysfunction (9). The new model system should now enable *in vitro* studies of how chronically stimulated CD8 T cells progress towards dysfunction under extended antigenic stimulation, and for assessment of microenvironmental cues beyond TCR signaling alone in this process.

### Chronic TCR plus TGF**β**1 signaling establishes the transcriptomic signature of terminal exhaustion and impairs protein translation

Next, we sought to determine whether *in vitro* chronic antigen plus TGFβ1 stimulation establishes the transcriptional circuits underlying T cell exhaustion *in vivo*. To investigate transcriptomic changes, we performed RNA sequencing on antigen-experienced P14 cells that were FACS-purified on day 7 (“Effector” stage) or day 19 (“Acute-2d”, “Acute-7d”, and “T_Dysf_”) (**Supplemental Figure 5A**). First, we used principal component analysis (PCA) to compare transcriptional profiles of these cells to *in vivo*-differentiated CD8 T cells, including memory and exhausted CD8 T cells from acute or chronic LCMV infections (28, 36). T_Dysf_ cells underwent distinct transcriptional programming compared to acutely stimulated or effector-like P14 cells, clustering closely with the terminally exhausted subset of TEX cells from chronic LCMV infection (28) (**Figure 3A, Supplemental Figure 5B**). In contrast, the transcriptomes of acutely stimulated P14 cells (Acute-2d and Acute-7d) closely resembled those of memory CD8 T cells, including central memory “TCM” and effector memory “TEM” cell subsets from acute LCMV infection (36), as well as progenitor TEX cells from chronic infection (28) (**Figure 3A, Supplemental Figure 5B**). These results demonstrate that prolonged TCR and TGFβ1 stimulation accurately models the transcriptional landscape of terminal T cell exhaustion.

**Figure 3.**
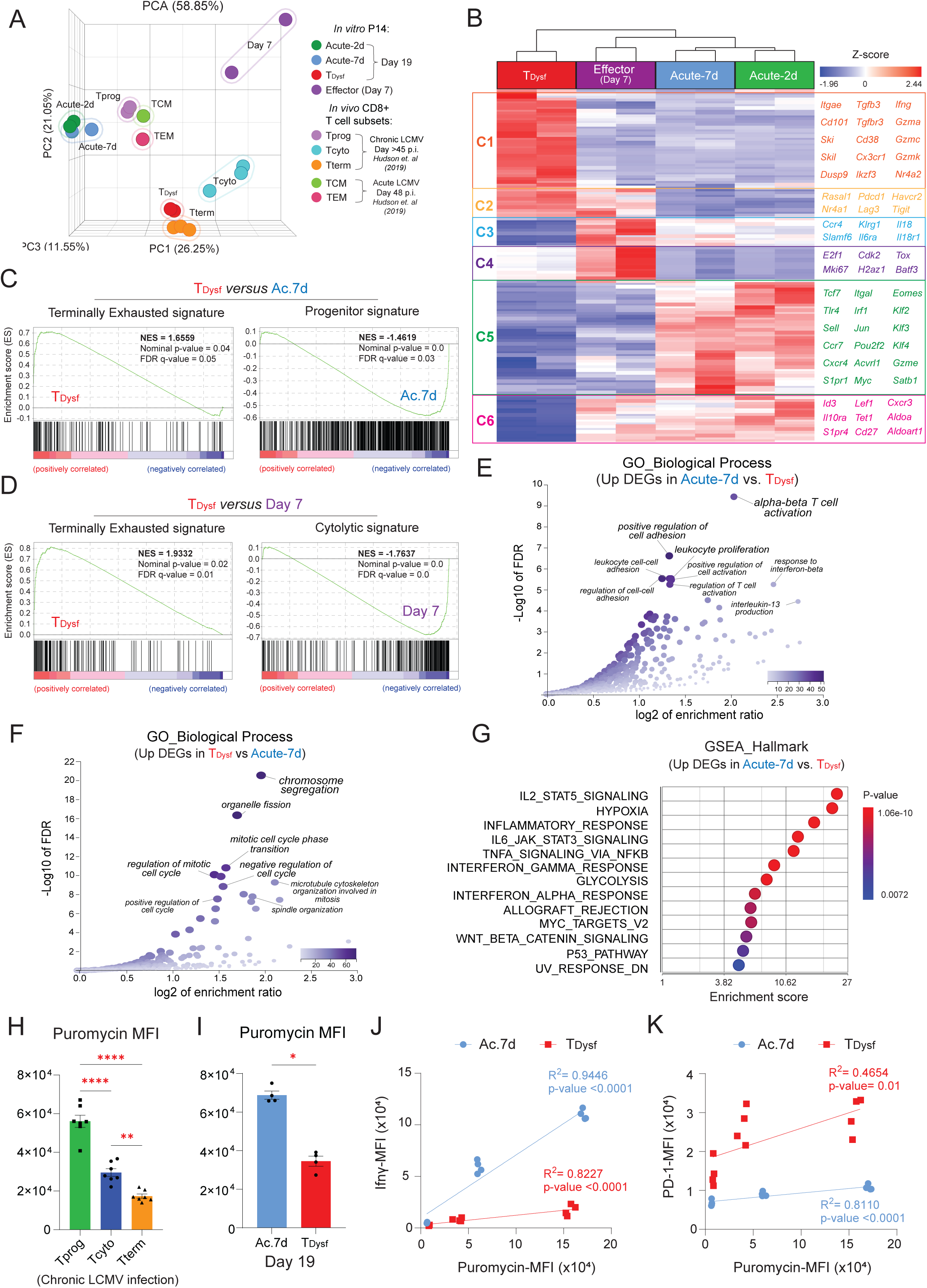
Chronic TCR plus TGFβ1 signaling establishes the terminal exhaustion transcriptome and impairs protein translation. (**A**) Principal component analysis (PCA) plot of RNA-sequencing data for the indicated antigen-stimulated P14 cell conditions isolated on either day 7 (Effector) or day 19 (all other conditions) of the *in vitro* model described in Figure 1A and **2A**: “T_Dysf_” (Chronic GP33 + TGFβ1), “Acute-7d” (GP33 stimulation for 7 days followed by rest), and “Acute-2d” (activation only followed by rest). P14 signatures were compared to published transcriptional signatures of exhausted PD-1+CD8+ T cell subsets on day >45 of chronic LCMV infection (Progenitor “Tprog” CD101-Tim3-, Cytolytic “Tcyto” CD101-Tim3+, Terminally Exhausted “Tterm” CD101+Tim3+) from Hudson et. al (28), and memory CD8 T cell subsets on day 48 post-acute LCMV infection (Central Memory “TCM”, Effector Memory “TEM”) from Hudson et. al (36). (**B**) Heatmap showing differentially expressed genes (DEGs) across the four P14 conditions. DEGs were plotted based on relative Z-score of normalized RNA counts and organized into clusters based on expression patterns. (**C**) Gene set enrichment analysis (GSEA) plots for DEGs upregulated in T_Dysf_ *versus* Acute-7d, or (**D**) T_Dysf_ *versus* Day 7 Effector in comparison to published signatures for TEX subsets on day >45 of chronic LCMV infection (28). (**E**) Gene ontology (GO) enrichment plot of biological processes for DEGs upregulated in Acute-7d *versus* T_Dysf_, or (**F**) upregulated in T_Dysf_ *versus* Acute-7d. (**G**) GSEA score plot compared to Hallmark gene signature for DEGs upregulated in Acute-7d *versus* T_Dysf_. (**H**) Summary bar graph for intracellular puromycin levels (gMFI) in TEX subsets isolated from LCMV clone 13-infected mice on day 21 post-infection, or (**I**) *in vitro*-stimulated P14 cells on day 19 after GP33 peptide rechallenge. (**J**) Summary plots showing intracellular levels of puromycin versus Ifnγ gMFI, or (**K**) PD-1 gMFI in P14 cells on day 19 after GP33 peptide rechallenge, with linear regression values listed. *N*=2 biological replicates for RNA-seq, or *n*□=□4-6 biological replicates for **H-K**, representative of two to three independent experiments. Statistical significance for (**A-B, E-G**) was determined by PCA, DESeq2 analysis or GSEA using Partek software, or (**C-D**) using UC San Diego and Broad Institute GSEA software. Comparisons in (**H**) were determined by One-way ANOVA, (**I**) Mann-Whitney U test (unpaired, two sided), or (**J-K**) simple linear regression. Adjusted P value \**P<0.05*, \*\**P<0.01*, \*\*\**P*□*<*□*0.001*, \*\*\*\**P*□*<*□*0.0001*. Error bars indicate meanL±LSEM.

T_Dysf_ cells, in particular, showed significant upregulation of exhaustion-associated genes, including surface inhibitory receptors shared with Effector cells (*Pdcd1, Havcr2, Tigit, Lag3*), as well as surface markers and transcription factors (TFs) linked to terminal exhaustion (*Cd101, Cd38, Tox, Nr4a1/2*) (**Figure 3B, Supplemental Figure 5C**). T_Dysf_ cells also upregulated the *Itgae* transcript encoding the integrin CD103, which is known to be induced by TGFβ1 signaling (37), as well as several negative regulators of TGFβ1 signaling (*Smurf1, Ski, Skil, Pmepa1*) (38). Notably, T_Dysf_ cells retained RNA expression of some effector molecules (*Ifng, Gzma, Gzmb, Gzmk*) compared to acutely stimulated cells (**Supplemental Figure 5C**). The discordant expression patterns of *Ifng* transcripts and protein in T_Dysf_ cells may be due to post-transcriptional regulation, impaired protein synthesis, or the chronic active stimulation conditions under which they were isolated, in contrast to acutely stimulated cells, which were resting before transcriptional analysis. Conversely, acutely stimulated P14 cells maintained higher expression levels of several stemness and memory-associated genes (e.g., *Sell, Ccr7, Tcf7, Lef1, Cd27*), along with genes linked to T cell migration and chemotaxis (e.g., *S1pr1, Klf2, Cxcr3, Cxcr4, S1pr4*) (**Figure 3B**, **Supplemental Figure 5C**).

To further determine whether the upregulated differentially expressed genes (DEGs) within T_Dysf_ cells recapitulate the transcriptional signature of exhausted T cells *in vivo*, we performed gene set enrichment analysis (GSEA) against the transcriptional signatures of progenitor, cytolytic and terminally exhausted TEX subsets from chronic LCMV infection in mice (28). Importantly, T_Dysf_ cells were significantly enriched for the terminally exhausted signature. In contrast, Acute-2d and Acute-7d cells closely resembled the progenitor subset signature, while the Effector-Day 7 P14 cells were enriched for the cytolytic subset signature (**Figure 3C-D**, **Supplemental Figure 5D**). Additionally, in comparison to single-cell transcriptional signatures of human TILs isolated from 21 types of cancer (33), we found that the T_Dysf_ cells were most enriched for TEX subsets, including KIR+TXK+ NK-like and terminal Tex TILs (**Supplemental Figure 5E**). Alternatively, the upregulated genes in Acute-7d P14 cells were enriched for functional subsets of TILs, most closely resembling the TEMRA transcriptional signature (**Supplemental Figure 5F**). To gain insights into the biological functions of upregulated genes in Acute-7d *versus* T_Dysf_ cells, we performed enrichment analysis using Gene Ontology biological processes and Hallmark GSEA, which revealed significant enrichment of pathways associated with T cell activation, proliferation, adhesion, IL-2/TNF/IFNG cytokine signaling, and glycolysis in the acutely stimulated T cells (**Figure 3E, G**). However, upregulated genes in T_Dysf_ were mainly enriched for cell cycle-related processes, E2F targets, and G2M checkpoint hallmarks (**Figure 3F, Supplemental Figure 5G-I**), previously linked with dysfunctional CD8 T cells (9, 39). Overall, our *in vitro*-generated T_Dysf_ cells closely recapitulate the transcriptional signature of terminally exhausted CD8 T cells in chronic infection and cancer.

In addition to transcriptional reprogramming, exhausted CD8 T cells also lose their capacity for protein translation during metabolic stress (40). To evaluate protein translational activity in TEX subsets from chronic LCMV infected mice, we performed a puromycin incorporation assay, in which puromycin, an aminoacyl-tRNA analog, is incorporated into nascent polypeptide chains during active protein synthesis (41). We observed a significant decrease of puromycin incorporation within terminally exhausted virus-specific CD8 T cells compared to the progenitor or cytolytic TEX subsets (**Figure 3H**), indicating impaired protein translation activity as CD8 T cells progress towards terminal exhaustion *in vivo*. Similarly, during *in vitro* GP33 peptide restimulation on day 19, T_Dysf_ cells showed significantly reduced incorporation of puromycin compared to Acute-7d cells (**Figure 3I, Supplemental Figure 5J**). Importantly, puromycin incorporation was positively correlated with expression of Ifnγ or PD-1 proteins in both Acute-7d and T_Dysf_ cells (**Figure 3J-K**). Together, these findings demonstrate that *in vitro*-generated T_Dysf_ cells acquire the transcriptional signature characteristic of terminally exhausted CD8 T cells in mice or humans, along with impairment in protein translation, which may explain the discordant RNA and protein expression patterns observed in TEX cells (42).

### Stable CD8 T cell dysfunction requires prolonged stimulation exceeding two weeks

The hallmark of terminal CD8 T cell exhaustion is the stability of the dysfunctional state (3, 10). To determine whether T_Dysf_ cells adopt this irreversible phenotype, we performed recovery experiments in which chronically stimulated P14 cells were withdrawn from antigenic stimulation and TGFβ1 signaling at different timepoints and allowed to rest under homeostatic conditions (**Figure 4A**). We first tested whether 12 days of chronic stimulation was sufficient to establish stable dysfunction by terminating GP33 and TGFβ1 exposure on day 12, followed by a 7 day-rest period until day 19 (“T_Dysf_-12d” condition). Importantly, T_Dysf_-12d cells demonstrated substantial functional recovery, with significant restoration of effector cytokine production and degranulation activity—reflected by increased frequencies of Ifnγ+ Tnf+, Ifnγ+ CD107a+, and Tnf+ CD107a+ cells—comparable to acutely stimulated cells (**Figure 4C-D, Supplemental Figure 6A**). This functional recovery was also accompanied by improved cell viability (**Figure 4B**). In contrast, T_Dysf_ cells rested from day 19 to day 26 failed to regain their effector function, maintaining significantly reduced cytokine production and degranulation capacity compared to acute control levels (**Figure 4C-D, Supplemental Figure 6A**). The viability of T_Dysf_ cells also significantly declined throughout the resting period (**Figure 4B**).

**Figure 4.**
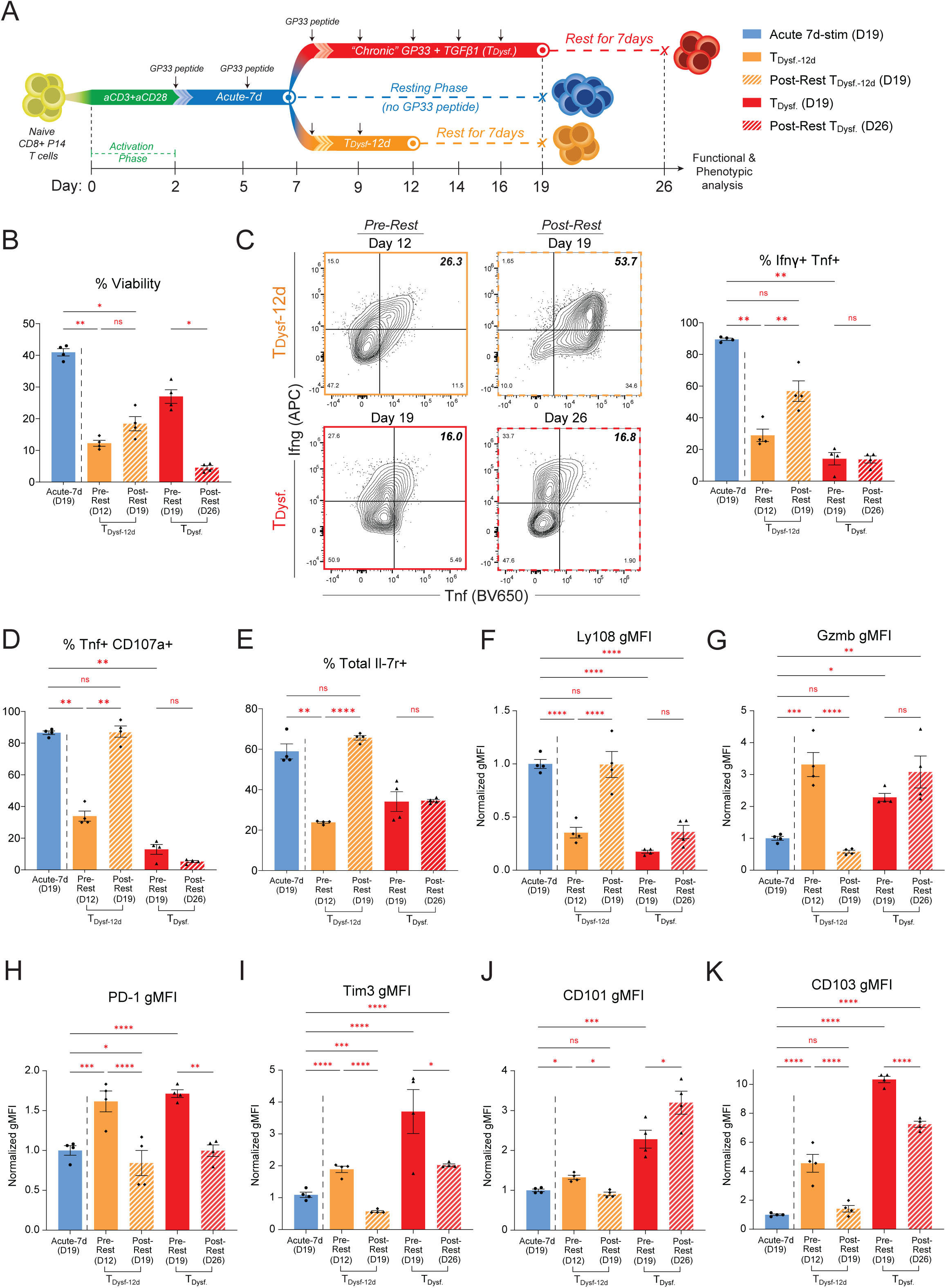
Stable CD8 T cell dysfunction requires prolonged stimulation exceeding two weeks. (**A**) Schematic showing resting phase of T_Dysf_ P14 cells under homeostatic conditions. Naïve P14 CD8+ T cells were stimulated with GP33 peptide and TGFβ1 for either 12 days (TDysf-12d) or 19 days (T_Dysf_), then rested in GP33 and TGFβ1-free medium for 7 days. (**B**) Bar graph depicting frequency of viable P14 cells comparing TDysf-12d pre-rest (day 12) and post-rest (day 19) vs TDysf. pre- and post-rest (day 19 and 26 respectively. (C-D) Functional assessment of polyfunctional cytokine production. Frequencies of (**C**) Ifnγ+Tnf+ and (**D**) Tnf+CD107a+ P14 cells following GP33 peptide rechallenge for P14 CD8+ T cells on day 12, 19, and 26 following resting for 7 days. (**E**) Bar graph showing frequency of Total Il-7r+ P14 cells. Expression level (normalized gMFI) of (**F**) Ly108, (**G**) Gzmb, (**H**) PD-1, (**I**) Tim3, (**J**) CD101, and (**K**) CD103 within P14 cells normalized to Acute-7d on day 19. All *n*□=□4 biological replicates, representative of two to three independent experiments. Adjusted P value \*\**P*□*<*□*0.01*, \*\*\**P*□*<*□*0.001*, \*\*\*\**P*□*<*□*0.0001*. Comparisons were determined by One-way ANOVA (**B-K**). Error barsLindicate meanL±LSEM.

Phenotypic analysis further distinguished reversible from stable dysfunction. T_Dysf_-12d cells exhibited significant downregulation of exhaustion markers during the rest period, including PD-1, Tim3, CD101, CD103, and granzyme B (**Figure 4G-K**), along with re-expression of memory-associated markers Ly108 and Il7r, though CD62L recovery remained modest (**Figure 4E-F**, **Supplemental Figure 6B-C**). This recovery profile suggests that 12 days of chronic stimulation induces a flexible, reversible dysfunctional state. In contrast, T_Dysf_ cells rested from day 19 retained high expression of terminal exhaustion markers and failed to re-express Il7r and Ly108, reflecting phenotypic rigidity and stable commitment to terminal exhaustion (**Figure 4E-F, Supplemental Figure 6B**). To further validate the stability of this state, we extended the recovery period to 11 days of resting (day 19 to day 30). Even after prolonged rest, T_Dysf_ cells failed to restore cytokine production, degranulation, or expression of memory markers, while exhaustion markers remained elevated (**Supplemental Figure 6D-L**). Together, these findings reveal a critical threshold in stimulation duration required for stable exhaustion programming. While less than two weeks of chronic stimulation results in reversible functional impairment, extending stimulation to ∼3 weeks establishes irreversible reprogramming characteristic of terminal dysfunction. The inability of terminally dysfunctional cells to regain function or memory potential even after prolonged rest supports the validity of this *in vitro* model as a faithful representation of irreversible T cell exhaustion observed *in vivo*.

### Dysfunctional P14 cells recapitulate the heterogeneity of exhausted CD8 T cells generated *in vivo*

T cell dysfunction is a gradual process that results in heterogeneous populations with different degrees of dysfunction. Indeed, on day 19 we identified distinct populations, with the predominant subset characterized as PD-1^hi^ Tim3+, and a minor subset of PD-1^int^ Tim3-cells, suggesting that such heterogeneity arises during chronic stimulation (**Figure 5A-B**). In contrast, resting Acute-7d P14 cells downregulate PD-1 and Tim-3 leading to the emergence of a PD-1^low^ Tim3-subset (**Figure 5A**). Across all these subsets, acutely stimulated P14 cells maintained significantly higher expression levels of effector cytokines and memory markers, such as Ifnγ, Tnf, CD107a, Ly108, Il7r, and CD62L, compared to T_Dysf_ P14 cells (**Supplemental Figure 7B-D, H**). Remarkably, the PD-1^hi^ Tim3+ subset within T_Dysf_ cells exhibited profound reductions in the expression of Perforin and the proliferation marker Ki67 (**Supplemental Figure 7F-G**), while also maintaining a significant increase in the expression of TOX (43) (**Supplemental Figure 7E**), further suggesting their terminally dysfunctional state.

**Figure 5.**
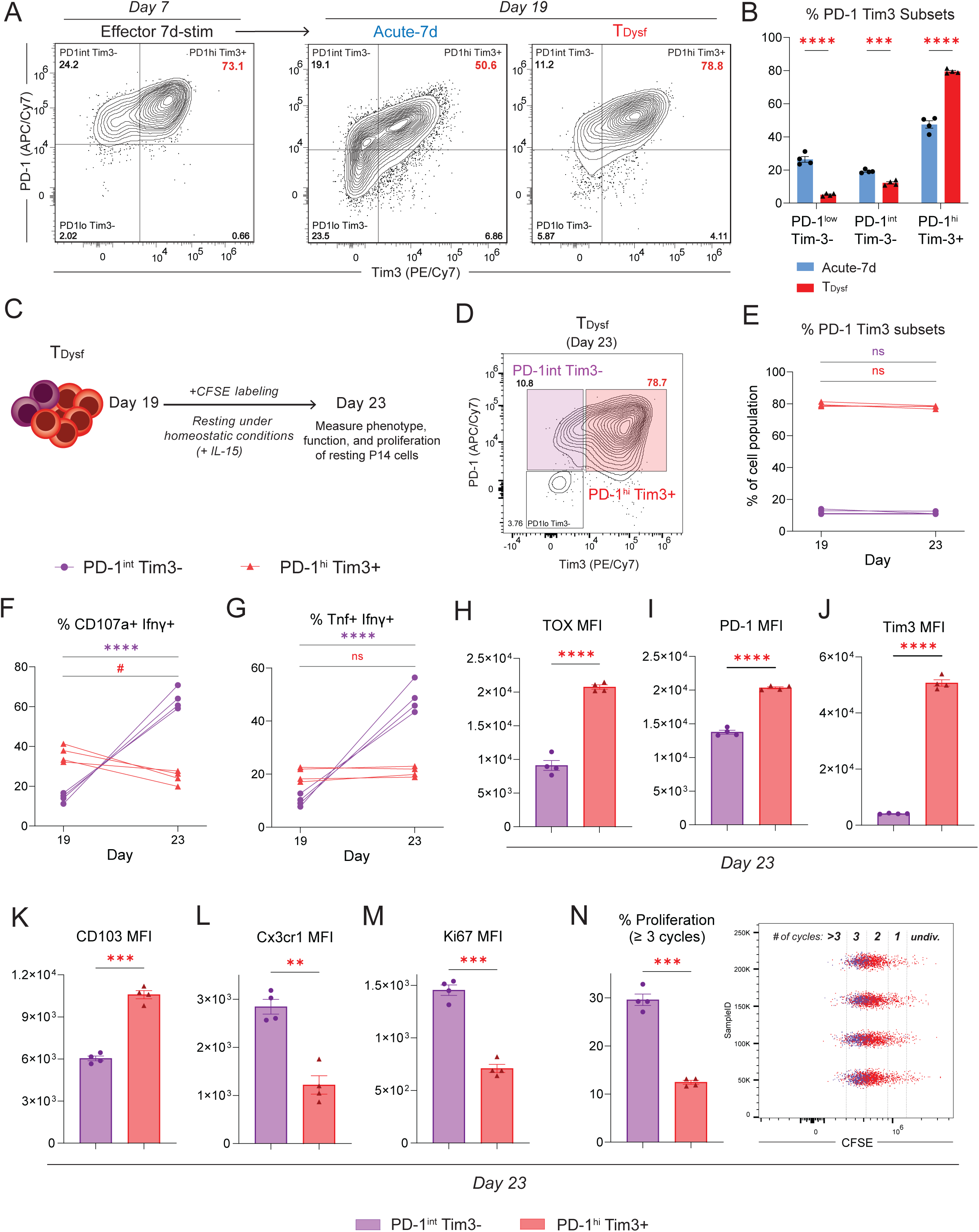
Dysfunctional CD8 T cells recapitulate the heterogeneity of exhausted CD8 T cells generated *in vivo*. (**A**) Representative FACS plots of PD-1 and Tim3 expression for P14 CD8+ T cells on both day 7 (Effector 7d-stim) and day 19 (Acute-7d or Chronic GP33+TGFβ1: T_Dysf_) after rechallenging with GP33 peptide. (**B**) Summary bar graph showing % of PD-1 and Tim3 expressing subsets on P14 CD8+ T cells on day 19. (**C**) Schematic for the resting phase of T_Dysf_ P14 cells under homeostatic conditions. (**D**) Representative FACS plot of PD-1 and Tim3 expression for P14 CD8+ T cells on day 23 following resting for 4 days. (**E**) Longitudinal tracking on day 19 and 23 of the frequency of PD1^int^ Tim3^−^ (purple) and PD1^hi^ Tim3+ (red) P14 cells, (**F**) frequency of CD107a+ Ifnγ+ P14 cells, and (**G**) frequency of Tnf+ Ifnγ+ P14 cells within PD1^int^ Tim3^−^ and PD1^hi^ Tim3+ subsets. (**H**) Expression levels (gMFI) of TOX, (**I**) PD-1, (**J**) Tim3, (**K**) CD103, (**L**) Cx3cr1, and (**M**) Ki67 within PD1^int^ Tim3^−^ and PD1^hi^ Tim3^+^ subsets on day 23. (**N**) Bar graph showing frequencies of divided P14 cells with > 3 proliferation cycles by CFSE staining. All *n*□=□4 biological replicates, representative of two to three independent experiments. Adjusted P value #*P<0.05* for PD-1^hi^ Tim3+ cells (**F**), \**P<0.05*, \*\**P*□*<*□*0.01*, \*\*\**P*□*<*□*0.001*, \*\*\*\**P*□*<*□*0.0001*. Comparisons were determined by One-way ANOVA (**B**), Two-way ANOVA (**E-G**), or Mann-Whitney U test (unpaired, two sided) (**H-N**). Error bars indicate meanL±LSEM.

To determine whether these T_Dysf_ P14 subsets have acquired a stable dysfunctional program, cells were CFSE-labeled and rested under homeostatic conditions for four days (day 19-23, **Figure 5C**). Importantly, the PD-1^hi^ Tim3+ subset maintained a severe dysfunctional state while retaining high expression levels of inhibitory receptors and the transcription factor, TOX (**Figure 5D-J, Supplemental Figure 7I-L**). In contrast, PD-1^int^ Tim3-cells demonstrated a remarkable recovery of polyfunctionality, associated with higher expression of Cx3cr1, which is a key marker of the cytolytic subset of TEX cells, compared to the PD-1^hi^ Tim3+ subset that maintained higher expression of CD103 (**Figure 5K-L**). Furthermore, PD-1^int^ Tim3-cells showed higher homeostatic proliferation potential with higher proportions undergoing at least 3 rounds of division during the resting phase (**Figure 5M-N**). Together, these results reveal that chronic stimulation over 3 weeks establishes a stable dysfunctional state within CD8 T cells, with the major PD-1^hi^ Tim3+ subset exhibiting features of terminal dysfunction.

### Terminal dysfunction of *in vitro* chronically stimulated CD8 T cells is stabilized by exhaustion-specific epigenetic programs

T cell exhaustion is marked by distinct epigenetic changes that lock terminally exhausted CD8 T cells into a stable dysfunctional state (3, 7–14). This epigenetic remodeling is characterized by progressive loss of chromatin accessibility at effector function, stemness and memory-associated genes, while maintaining open chromatin at exhaustion-related loci, such as inhibitory receptors (11, 23, 44). To define epigenetic changes within our *in vitro*-generated P14 cells, we performed the Assay for Transposase-Accessible Chromatin using sequencing (ATAC-seq) on Acute-7d and T_Dysf_ (PD-1+ Tim3+) P14 cells isolated on day 19 (**Figure 6A**). T_Dysf_ cells acquired a distinct open chromatin landscape, with over 24,600 differentially open chromatin regions (OCRs; *P-value <0.01*, fold change ≥2), including ∼13,900 regions with reduced accessibility and ∼10,700 regions with increased accessibility relative to Acute-7d cells (**Figure 6B**). These OCRs were primarily located in intronic and intergenic regions, suggesting regulatory roles (**Supplemental Figure 8A-B**). Regions with increased accessibility in T_Dysf_ cells included genes involved in TGFβ1 signaling (e.g., *Tgfb1, Tgfbr2, Pmepa1, Skil, Smad9*), genes encoding inhibitory receptors (e.g., *Pdcd1, Cd200r2*), MAPK signaling-related molecules (e.g., *Dusp16*, *Map2k2, Map2k4*), and exhaustion-associated transcription factors (*Nr4a3, Stat3*, and *Irf8*) (45, 46) (**Figure 6B**). In contrast, regions with reduced accessibility included stemness and memory genes, such as *Ccr7, Sell, Slamf6, Cxcr5*, and *Il15ra*, and TCR signaling and effector differentiation genes (e.g., *Cd28, Cd44, Il2ra, Icos, Icosl*, *Itgal*, *Ifngr1* and *Ifngr2*) (2, 29, 47) (**Figure 6B**). Pathway enrichment analysis revealed that closed OCRs in T_Dysf_ cells were enriched for IL-2/STAT5, TCR signaling, interferon gamma and inflammatory response pathways, while open regions were enriched for PI3K/AKT/mTOR, Hedgehog and Notch signaling, and GTPase activity pathways (**Supplemental Fig.8C-F**). Together, these results demonstrate extensive epigenetic remodeling within T_Dysf_ cells, marked by reduced chromatin accessibility at effector and memory-associated genes and increased accessibility at exhaustion-linked loci, reflecting a stable, epigenetically reprogrammed dysfunctional state.

**Figure 6.**
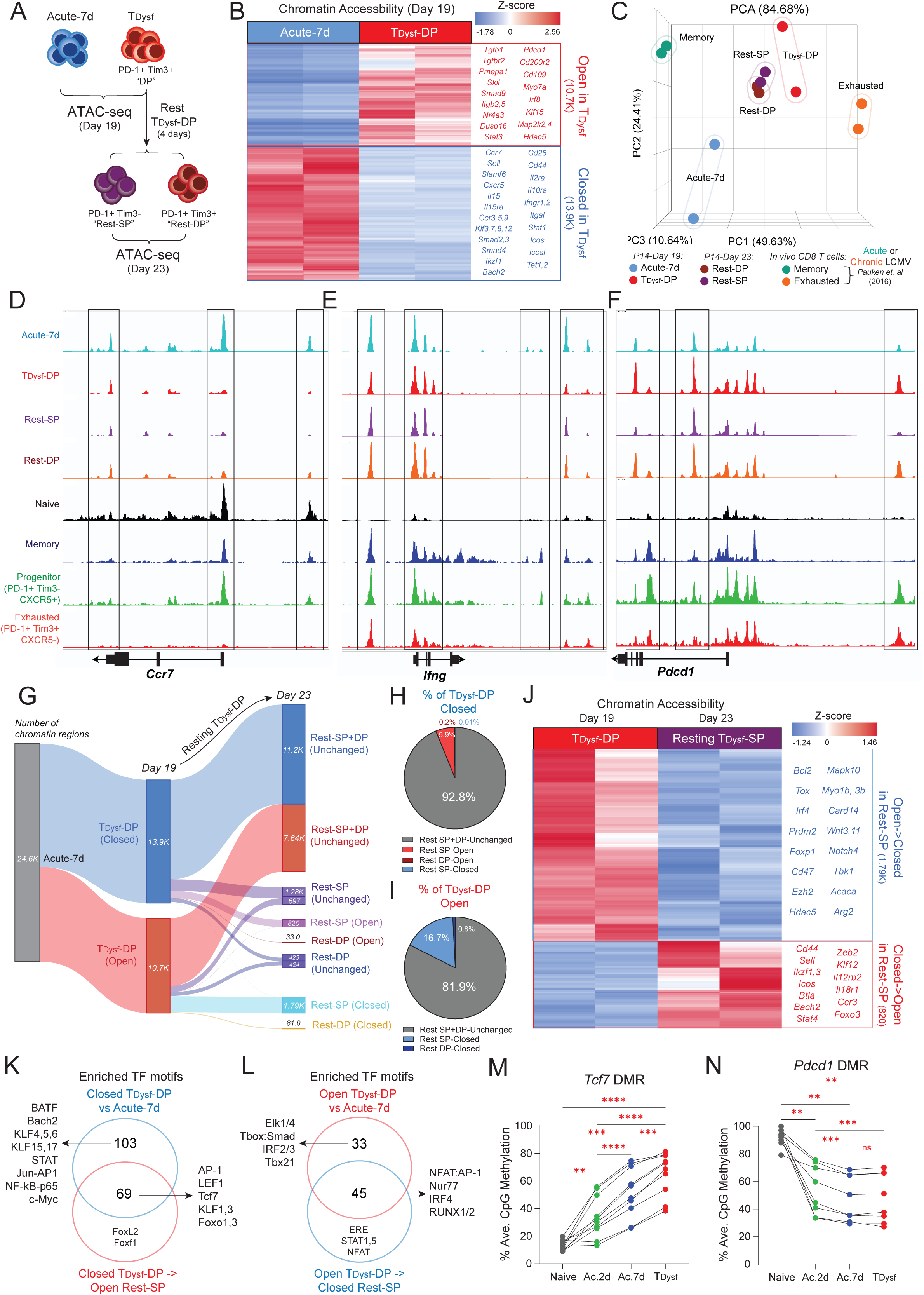
Terminal dysfunction of *in vitro*-generated CD8 T cells is stabilized by exhaustion-associated epigenetic programs. (**A**) Schematic for ATAC-sequencing of *in vitro* acutely or chronically stimulated P14 CD8 T cells isolated on Day 19 or Day 23 of the model described in Fig.2A and **5C**. For T_Dysf_ condition, cells were FACS-sorted on Day 19 of chronic stimulation as the PD-1+Tim3+ subset (double-positive, “T_Dysf_-DP”) and rested for 4 days until Day 23. Resting T_Dysf_-DP cells were sorted on Day 23 as either PD-1+Tim3-(single-positive, “Rest-SP”) or PD-1+Tim3+ (double-positive, “Rest-DP”). (**B**) Heatmap showing differentially open chromatin regions (OCRs) between Acute-7d and T_Dysf_-DP cells on Day 19 with example genes listed. Regions were plotted based on relative Z-score with red indicating more accessible or “open” regions, and blue indicating less accessible or “closed” regions. (**C**) Principal component analysis (PCA) plot comparing global chromatin accessibility profiles for *in vitro* Acute-7d or T_Dysf_ P14 subsets to published signatures for *in vivo* memory or exhausted CD8 T cells (23). (**D**) Representative IGV snapshots of mapped accessible chromatin peaks within *in vitro* P14 or *in vivo* naive, memory, progenitor or exhausted TEX subsets (44) at gene loci for *Ccr7*, (**E**) *Ifng*, and (**F**) *Pdcd1*. (**G**) Alluvial plot tracking the number of chromatin regions from Acute-7d cells that became more Open/Closed in T_Dysf_-DP on Day 19, and regions from T_Dysf_-DP cells that remain Unchanged or became more Open/Closed in Resting T_Dysf_ cells on Day 23. (**H**) Pie chart showing frequency of Closed OCRs in T_Dysf_-DP Day 19 cells that remain Unchanged or became more Open/Closed in Resting T_Dysf_ cells, and (**I**) Open OCRs in T_Dysf_-DP Day 19 cells that remain Unchanged or became more Closed in Resting T_Dysf_ cells. (**J**) Heatmap showing differentially OCRs between T_Dysf_-DP cells on Day 19 and Resting T_Dysf_-SP cells on Day 23 with example genes listed. (**K**) Venn diagram comparing transcription factor (TF) motifs that were enriched in Closed OCRs in T_Dysf_-DP Day 19 cells and in regions that became Open in Resting SP cells, and (**L**) Open OCRs in T_Dysf_-DP Day 19 cells and regions that became Closed in Resting SP cells with example TFs listed. (**M**) % methylation of individual CpG sites at key differentially methylated regions (DMRs) at *Tcf7*, and (**N**) *Pdcd1* loci within naive or *in vitro*-generated P14 cells on day 19 (“Acute-2d”, “Acute-7d”, or “T_Dysf_”). *N*=2 biological replicates for ATAC-seq and DNA methylation sequencing, representative of two to three independent experiments. Comparisons (**M-N**) were determined by repeated-measures one-Way ANOVA analysis with Tukey’s multiple comparisons test. Adjusted P value ** P<0.01, \*\*\**P<0.001*, \*\*\*\**P*□*<*□*0.0001*.

To determine whether the epigenetic landscape of T_Dysf_ cells mirrors exhaustion-specific programming observed *in vivo*, we compared chromatin accessibility profiles of T_Dysf_ and Acute-7d P14 cells with the published ATAC-seq data of virus-specific CD8 T cells isolated during acute and chronic LCMV infection (23). PCA analysis of the top 5,000 most variable peaks revealed that T_Dysf_ cells closely resembled terminally exhausted TEX cells, while Acute-7d cells clustered closer to memory CD8 T cells (**Figure 6C**). Importantly, differentially accessible OCRs between T_Dysf_ and Acute-7d cells reflected patterns observed in terminally exhausted (PD-1+Tim3+CXCR5-) *versus* progenitor (PD-1+Tim3-CXCR5+) TEX cells or memory CD8 T cells (44). For instance, T_Dysf_ cells exhibited reduced accessibility at memory and progenitor TEX-associated genes involved in stemness and effector function (e.g., *Ccr7, Ifng, Tnf, Cxcr5, Sell, Slamf6, Il7r, Tcf7*), and increased accessibility at terminal exhaustion-associated genes (e.g., *Pdcd1, Havcr2, Cd244a*) (**Figure 6D-F**, **Supplemental Figure 8I-Q**). These findings indicate that T_Dysf_ cells undergo exhaustion-specific chromatin remodeling consistent with terminally exhausted CD8 T cells *in vivo*, whereas Acute-7d cells retain a memory-like open chromatin landscape.

Next, we examined whether the dysfunction-associated chromatin accessibility landscape remains stable after T_Dysf_ cells are withdrawn from chronic stimulation. To define chromatin accessibility dynamics upon rest (**Figure 5**), we performed ATAC-seq on sorted PD-1^hi^ Tim3+ cells (“double-positive”, Rest-DP) and PD-1^int^ Tim3-(“single-positive”, Rest-SP) P14 cells isolated on day 23 following withdrawal from chronic antigen and TGFβ1 signaling (**Figure 5C-D**, **Figure 6A**). Both Rest-DP and Rest-SP populations largely retained the chromatin accessibility profiles of T_Dysf_ cells and terminally exhausted CD8 T cells (**Figure 6C-F**, **Supplemental Figure 8I-Q**). Notably, ∼92.8% of T_Dysf_-specific closed OCRs and ∼81.9% of open OCRs remained unchanged in the rested populations (**Figure 6G-I**), indicating strong epigenetic stability. However, the Rest-SP population showed partial reversal of a small fraction of T_Dysf_ chromatin accessibility programs. Specifically, ∼5.9% (820 OCRs) of previously closed regions regained accessibility, including genes such as *Cd44*, *Sell*, *Btla* and *Icos*, transcription factors (*Izkf1*, *Ikzf3*, *Bach2*, *Stat4*, *Zeb2*, *Klf12*, and *Foxo3*), and cytokine receptors (*Il12rb2*, *Il18r1*) (**Figure 6G-J**). Conversely, ∼16.7% (∼1790 OCRs) of T_Dysf_-associated open regions became less accessible, including peaks at TFs (e.g., *Bcl2*, *Tox*, *Irf4*, *Prdm2*, and *Foxp1*), histone-modifying enzymes (e.g., *Ezh2* and *Hdac5*), and signaling molecules (e.g., *Mapk10, Wnt3, Wnt11, Notch4, Tbk1*) (**Figure 6G-J**). Pathway enrichment analysis of regions that recovered accessibility in Rest-SP cells included IL-2 and TCR signaling pathways, while regions that lost accessibility in resting cells were enriched for GTPase activity and MAPK signaling pathways (**Supplemental Figure 8G-H**). Together, these findings demonstrate that T_Dysf_ cells retain a largely stable, epigenetically “scarred” chromatin state after removal of chronic signals—particularly in PD-1+ Tim-3+ cell population—closely resembling the fixed epigenetic profile of terminally exhausted CD8 T cells *in vivo* (11, 23, 44).

To gain insights into molecular mechanisms underlying the stability of dysfunction-associated epigenetic programs, we performed TF motif enrichment analysis on T_Dysf_-specific OCRs that remained unchanged following resting. Stable closed OCRs were enriched for motifs of TFs involved in effector and memory programs, including BATF, Bach2, AP1, NF-κB, c-Myc, KLF, and STAT family members. Importantly, Rest-SP cells showed partial recovery of chromatin accessibility at peaks enriched for Tcf7, LEF1, KLF1/3, and Foxo1/3 motifs (**Figure 6K**), suggesting limited re-engagement of memory-related transcriptional regulators. Conversely, stable open OCRs in Rest-SP and Rest-DP cells were enriched for motifs associated with known and potential exhaustion-linked TFs, including Elk1/4, Smad, IRF2/3, and Tbx21 motifs. Meanwhile, Rest-SP cells exhibited reduced chromatin accessibility at OCRs containing NFAT, Nur77, IRF4, RUNX1/2, and STAT1/5 motifs, suggesting re-wiring of specific TCR and cytokine-dependent signaling pathways in this subset that partially restores function following the resting period (**Figure 6L**). Overall, these findings reveal that specific TF networks enforce the stability of the dysfunctional epigenetic state in T_Dysf_ cells by maintaining accessibility at exhaustion-related motifs and closing memory- and effector-regulatory regions, highlighting a fixed transcriptional remodeling that limits recovery even after chronic stimulation is withdrawn.

Our previous work has demonstrated that de novo DNA methylation drives epigenetic silencing of memory and effector programs within TEX cells (8, 9). Additionally, specific DNA demethylation changes at the *Pdcd1* locus promote PD-1 upregulation in TEX cells, beginning in the effector phase of CD8 T cell differentiation (26, 48). To determine whether T_Dysf_ cells acquire distinct DNA methylation programs associated with stable exhaustion, we performed targeted DNA methylation analysis at some key differentially methylated genomic regions (DMRs) (8) (**Supplemental Figure 9A-C**). Our analysis revealed that T_Dysf_ cells acquired substantial de novo DNA methylation at both *Tcf7* and *Ccr7* loci, which are associated with significant downregulation in both Tcf1—a key transcription factor regulating stemness and memory programming—and Ccr7, which is an important chemokine receptor regulating lymphoid tissue-homing properties of memory T cells (**Figure 6M, Supplemental Figure 9D**). These de novo DNA methylation programs were associated with a significant downregulation of both Tcf1 and Ccr7 proteins by T_Dysf_ cells, while acutely stimulated cells were able to recover higher expression levels of both molecules on day 19 (**Supplemental Figure 9E-I**). In contrast, the *Pdcd1* DMR underwent significant DNA demethylation in both Acute and T_Dysf_ P14 cells (**Figure 6N**), consistent with the reported DNA demethylation at *Pdcd1* during the acute phase of CD8 T cell differentiation (26). However, PD-1 was augmented even further on T_Dysf_ relative to acutely stimulated cells on day 19, likely driven by chronic antigen stimulation (26, 27, 49) (**Supplemental Figure 9E-G**). We conclude that acquisition of *de novo* DNA methylation limits the ability of dysfunctional P14 cells to retain key stemness and memory features, and parallels cell-intrinsic defects in these critical programs in terminally exhausted CD8 T cells *in vivo*.

### Dysfunction of *in vitro*-generated T_Dysf_ cells is irreversible during *in vivo* rechallenge

Finally, we assessed the *in vivo* recall responses of our *in vitro*-generated T_Dysf_ cells, as a key hallmark of T cell exhaustion is the compromised ability to respond following antigen rechallenge (11, 24). We sorted congenically distinct Thy1.1+ P14 cells from T_Dysf_ or Acute-7d groups on day 19, followed by adoptive transfer into naive C57BL/6 animals. One day later, we challenged the mice with acute LCMV infection and measured the recall responses of P14 cells after 6-7 days of infection **(Figure 7A)**. While Acute-7d P14 cells underwent extensive antigen-dependent proliferation, T_Dysf_ cells showed compromised proliferation responses during infection **(Figure 7B**), as well as sustained downregulation of Ly108 **(Figure 7C).** In addition, after *ex vivo* GP33 peptide restimulation of splenocytes, T_Dysf_ cells showed poor effector functions, indicated by significantly lower expression of effector molecules (Ifnγ and Perforin) compared to Acute-7d P14 cells, which maintained heightened polyfunctionality **(Figure 7D-E)**. T_Dysf_ cells displayed phenotypic characteristics of terminally exhausted T cells, including elevated levels of exhaustion-associated molecules, such as Tox and Gzmb compared to Acute-7d P14 cells within the spleen, liver, and lungs **(Figure 7F-G, Supplemental Figure A-B)**. To test whether resting T_Dysf_ cells improves their recall capacity, we adoptively transferred rested T_Dysf_ P14 cells into C57BL/6 mice after a 7-day rest period and then challenged them with acute LCMV infection (**Figure 7H**). Despite the rest, T_Dysf_ cells showed impaired proliferation and reduced expression of memory- and stemness-associated markers, such as Ly108 and Il7r (**Figure 7I-J, Supplemental Figure 10C**). They also exhibited diminshed effector function (e.g. Ifny, Tnf, and CD107a) (**Figure 7K)** and retained high expression of exhaustion-related markers, including PD-1, Tim- 3, Gzmb and Tox (**Figure 7L-M, Supplemental Figure 10D-F)**. Taken together, these findings demonstrate that ∼3 weeks of chronic stimulation *in vitro* drives CD8 T cells into a stable, terminally dysfunctional program that closely mirrors *in vivo* terminal exhaustion, including severely impaired ability to recall effector functions.

**Figure 7.**
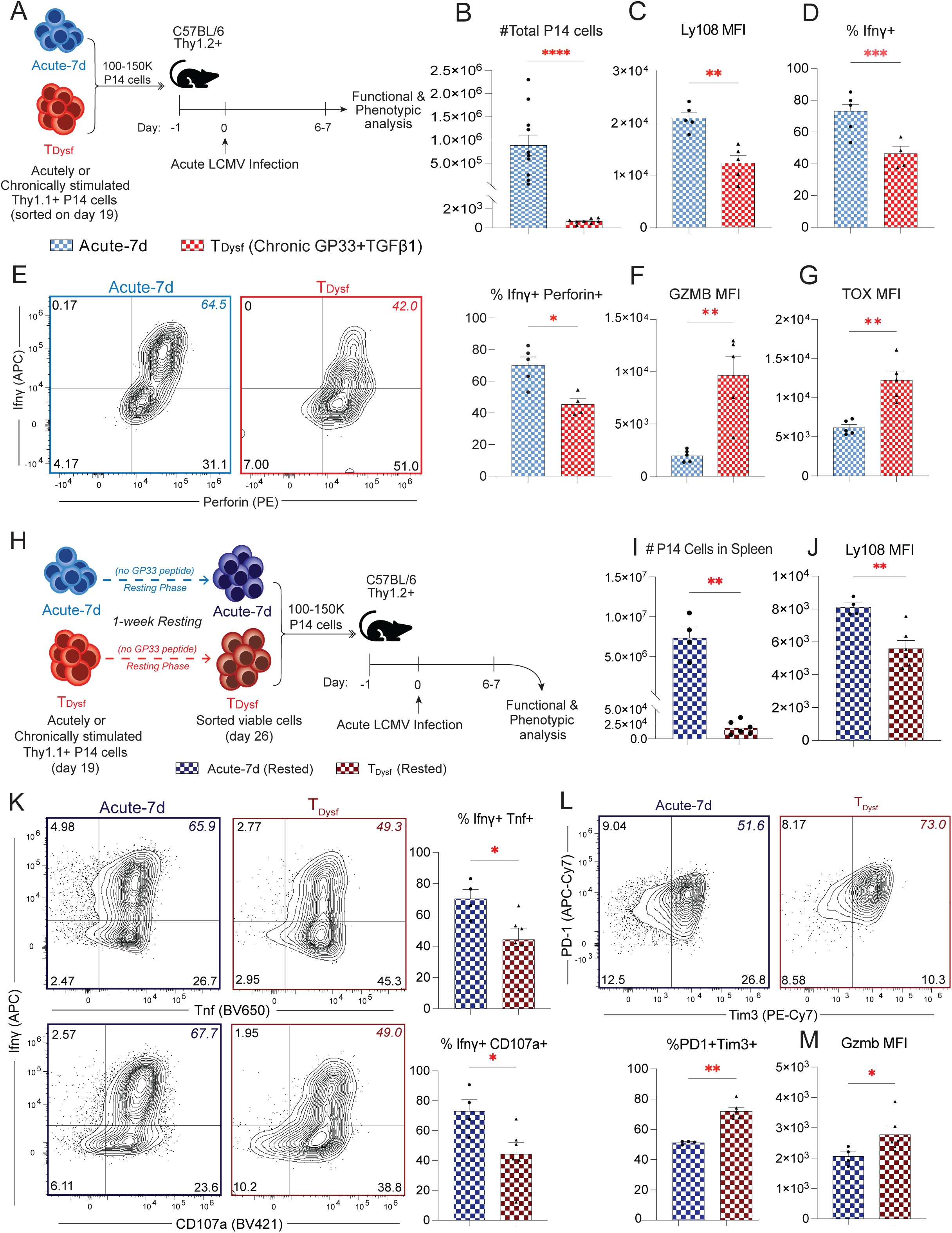
Dysfunction of *in vitro*-generated T_Dysf_ cells is irreversible during *in vivo* rechallenge. **(A)** Schematic for adoptive transfer of 100,000-150,000 “Acute-7d” (blue) or “T_Dysf_” (red) Thy1.1+ P14 CD8+ T cells generated *in vitro* into Thy1.2+ C57BL/6 mice on day -1, followed by acute LCMV Armstrong infection on day 0. Functional and phenotypic analysis of adoptively transferred cells was performed on days 6 or 7 post-infection. **(B)** Total number of P14 cells from the spleens, livers, and lungs of mice harvested on day 6 of acute LCMV infection. **(C)** Bar graph showing the expression level (gMFI) of Ly108 in P14 cells isolated from the spleens. **(D)** Bar graph showing the % of Ifnγ+ P14 cells after *ex vivo* 5-hr GP33 peptide rechallenge. (**E**) FACs plots showing Ifnγ and Perforin expression and summary bar graph showing % of Ifnγ+ Perforin+ P14 cells isolated from the spleens. The 7dAc plot is representative of one 7dAc sample, while the T_Dysf_ plot shows data from all T_Dysf_ samples. **(F)** Bar graph showing the expression level (gMFI) of Gzmb and **(G)** TOX in P14 cells isolated from the spleens. **(H)** Schematic showing experimental design used to generate data for panels **I**-**M**. *In vitro* generated 7-day Acute and T_Dysf_ P14 T cells were rested from day 19 to day 26 (7 days), followed by adoptive transfer of 100,000-150,000 “Rested Acute-7d” (dark blue) or “Rested T_Dysf_” (dark red) Thy1.1+ P14 CD8+ T cells into Thy1.2+ C57BL/6 mice on day -1, followed by acute LCMV Armstrong infection on day 0. Functional and phenotypic analysis of adoptively transferred cells was performed on days 6 or 7 post-infection. (**I**) Bar graph showing the total number of P14 cells isolated from the spleens. (**J**) Bar graph showing the expression level (gMFI) of Ly108 in P14 cells isolated from the spleens. (**K**) FACs plots showing Ifnγ and Tnf/ CD107a expression and summary bar graph showing % of Ifnγ+ Tnf+ as well as % of Ifnγ+ CD107a+ P14 cells isolated from the spleens. The 7dAc plot is representative of one 7dAc sample, while the T_Dysf_ plot shows data from all T_Dysf_ samples. (**L**) FACs plots showing PD-1 and Tim-3 expression and summary bar graph showing % of PD-1+ Tim-3+ P14 cells isolated from the spleens. The 7dAc plot is representative of one 7dAc sample, while the T_Dysf_ plot shows data from all T_Dysf_ samples. (**M**) Bar graph showing the expression level (gMFI) of Gzmb in P14 cells isolated from the spleens. For **B**, data were pooled from two independent experiments with *n*□=□3–5 biological replicates per group for each experiment. For all other panels *n*□=□4-6 biological replicates, representative of two to three independent experiments. Adjusted P value \**P<0.05*, \*\**P <0.01,* \*\*\**P*□*<*□*0.001*, \*\*\*\**P*□*<*□*0.0001*. Comparisons were determined by the Mann-Whitney U test (unpaired, two sided). Error bars indicate meanL±LSEM.

## Discussion

Epigenetic scarring of T cell exhaustion programs is a major cell-intrinsic barrier to T cell-based immunotherapies. Faithful model systems for T cell exhaustion are essential for understanding the individual and cooperative contributions of factors driving the epigenetic progression towards terminal exhaustion. While common animal models of T cell exhaustion, such as chronic LCMV infection or tumor models, are among the most reliable systems for these investigations, they present inherent complexities that make it challenging to dissect the precise contributions of individual factors to the epigenetic programming of TEX cells. Therefore, there is a major need to develop reliable reductionist approaches that enable precise control and isolation of individual signals, facilitating a deeper understanding of how specific factors influence the programming events underlying T cell exhaustion.

Recent studies have introduced new *in vitro* models of T cell dysfunction through repeated polyclonal anti-CD3 or antigen-specific stimulation of murine CD8 T cells (18–21). However, a key limitation has been the poor survival and extensive cell death of repeatedly TCR-stimulated CD8 T cells, preventing chronic stimulation from extending beyond 5-10 days (20). This restricts the ability to replicate *in vivo* settings, such as chronic viral infections, where CD8 T cells experience prolonged stimulation over ∼3 weeks before developing a terminally dysfunctional state (2, 24). Indeed, we found that ∼1-2 weeks of repeated stimulation induces some exhaustion-like features, marked by increased co-expression of inhibitory receptors; however, these conditions fail to establish epigenetically stable terminal dysfunction. Instead, after acute antigen stimulation is removed, CD8 T cells rapidly recover effector functions and memory programs. These findings align with the demonstrated plasticity of day 8-15 effector CD8 T cells, which can regain function and memory potential when removed from chronically LCMV-infected environments and transferred to antigen-free conditions (24). Thus, short-term antigen stimulation alone is insufficient for inducing terminal dysfunction; instead, it drives effector differentiation with preserved memory potential, mimicking CD8 T cell responses during acute viral infections, during which viruses are typically cleared within 1-2 weeks (50).

Here, we have developed a faithful animal cell model for terminal exhaustion by delivering chronic TGFβ1 signals between days 7 and 19 to chronically antigen-stimulated P14 CD8 T cells. This *in vitro* model builds on our previous work showing that chronic, post-effector TGFβ1 signaling is necessary for driving terminal dysfunction in chronically TCR-stimulated human CD8 T cells (9). Here, we demonstrate that prolonged TGFβ1 signals are critical for maintaining the survival of chronically antigen-stimulated P14 (T_Dysf_) cells, while also promoting an epigenetically stabilized, terminally dysfunctional state. Under these conditions, T_Dysf_ cells exhibited the full range of phenotypic, functional, transcriptional, epigenetic, and metabolic characteristics of terminal exhaustion. The stability of this terminal exhaustion program was validated by resting T_Dysf_ cells following antigen withdrawal, which revealed persistent epigenetic “scarring” of the dysfunctional state. A limited degree of epigenetic flexibility was observed only within the minor PD-1+ Tim-3-subset that emerged after resting T_Dysf_ cells. Furthermore, adoptive transfer studies of T_Dysf_ cells—both before and after resting—demonstrated that T_Dysf_ cells remained terminally dysfunctional, as they failed to regain effector functions or proliferate in response to acute LCMV infection.

The new, tractable model of T cell exhaustion described here offers several advantages for studying molecular mechanisms that underline terminal dysfunction. Specifically, the *ex vivo* culture model: a) enables tracking of the developmental trajectory of T cell exhaustion over extended periods, b) facilitates mechanistic studies of various signals from infected-tissue or tumor microenvironments, c) supports high-throughput molecular investigations of terminal T cell exhaustion improving both the technical and economic efficiency, compared to genetic or therapeutic screening studies performed *in vivo*, d) provides a more reliable tool for identifying key factors that stabilize the dysfunctional state in CD8 T cells, and e) offers physiologically relevant conditions for studying antigen-specific T cell responses by using antigen-dependent stimulation, rather than polyclonal, CD3-dependent stimulation. The T_Dysf_ model can be leveraged to design an effective discovery platform for the development of more effective immunotherapies aimed at restoring the function of severely dysfunctional T cells, ultimately enhancing treatment outcomes for patients with cancer or chronic infections.

## Materials and Methods

All research conducted in this study complies with the ethical regulations under the approved protocols by the Ohio State University Institutional Animal Care and Use Committee (IACUC #2019A00000055-R1) and the Institutional Biosafety Committee (IBC #2020R00000129).

### Animal housing conditions

All mice used in the LCMV infection experiments from this study were housed at the Ohio State University animal facilities under the IACUC-approved guidelines. This facility included an ambient temperature kept in a range of 68–76 °C, relative humidity 30–70%, and a 12-h dark/12-h light cycle (light 6 AM–6 PM). Mice were fed standard Teklad 7912 chow (Envigo).

### *In vitro* chronic stimulation of mouse P14 CD8 T cells

On Day 0 of the *in vitro* chronic stimulation model, CD8+ T cells were first isolated from the spleen of healthy, uninfected male or female P14 transgenic mice (at least 8 weeks old), in which all T cells are specific to the LCMV glycoprotein 33 (GP33) antigen. Spleens were harvested and processed by mashing over a 100 µm filter followed by red blood cell (RBC) lysis. Splenocytes were then enriched for total CD8+ T cells using the EasySep™ Mouse CD8+ T Cell Isolation Kit (StemCell Technologies), then stained for spectral cytometry to confirm purity, viability and GP33 tetramer specificity. The isolated CD8+ T cells were then activated by seeding into 6-well, tissue culture-treated flat-bottom plates previously coated with mouse purified anti-CD3 (5 µg/ml) and anti-CD28 (10 µg/ml) (BioLegend), at a density of 1-3 × 10^6^ CD8+ T cells per well in 3 ml of “complete” RPMI medium (containing 10% fetal bovine serum and 1X Penicillin/Streptomycin (Gibco) plus 200 IU/ml of recombinant human IL-2 (PeproTech) and 50 µM of 2-Mercaptoethanol (Millipore Sigma) to stabilize IL-2 activity and CD8+ T cell priming (51). Plates were placed in an incubator at 37 degrees Celsius and 5% CO_2_ for 2 days to allow T cell activation.

On Day 2 following incubation, the activated P14 CD8 T cells were pooled and centrifuged to remove activation medium. The cell pellet was resuspended in “T Cell Media” (complete RPMI medium containing 20 IU/ml of recombinant human IL-2 and 10 ng/ml of recombinant human IL-15 (PeproTech)). P14 cells were then seeded into tissue culture-treated 96-well U-bottom plates at a density of 1-2 e5 viable cells per well. For the 2d-stimulated condition, cells were seeded in T cell media without GP33 peptide. All other conditions (“Acute-7d” and “Chronic” conditions) were seeded in T Cell Media containing GP33 peptide at a final concentration of 0.3 µg/ml. On Day 7, “Acute-7d” cells were continued in T Cell Media without peptide throughout the remainder of the experiment. For the Chronic plus TGFβ1 “T_Dysf_” condition, recombinant human TGFβ1 (PeproTech) was added to the GP33 peptide-containing medium at a final concentration of 5 ng/ml beginning on Day 7. Anti-CD28 was omitted during the chronic stimulation phase to simulate the declining costimulatory environment present in chronic viral infections and tumors, where CD28 expression is progressively lost as TEX cells become terminally exhausted. Media was replenished every 2-3 days throughout the experiment with GP33 peptide and TGFβ1 until Day 19.

On Day 19 of chronic stimulation, P14 cells were collected for sorting of the viable CD8+ population (using Sony MA900 Cell Sorter). Sorted cells were either used immediately for adoptive transfer *in vivo*, for *in vitro* recovery experiments or ATAC-sequencing, or snap frozen at –80 °C to use for downstream RNA and DNA methylation sequencing analyses.

For “recovery” experiments, sorted P14 CD8 T cells from Day 12 or 19 of chronic stimulation were seeded in 96-well U-bottom plates at a density of 5 × 10^4^ cells per well, in complete RPMI medium containing 10 ng/ml of IL-15. Cells were cultured in this medium for 4-12 days, then used for functional analysis on Day 23, Day 26, and Day 30 of *in vitro* culture.

### Plate-Bound *in vitro* stimulation of mouse P14 CD8 T cells

Following a two-day activation phase with mouse purified anti-CD3 (5 µg/ml) and anti-CD28 (10 µg/ml) (BioLegend), activated P14 CD8 T cells were resuspended in complete RPMI medium containing 20 IU/ml of recombinant human IL-2 and 10 ng/ml of recombinant human IL-15 (PeproTech). P14 cells were then seeded into tissue culture-treated 96-well flat-bottom plates coated with anti-CD3 (5 µg/ml) at a density of 1-2e5 viable cells per well. Cells were transferred to freshly coated anti-CD3 plates every two days and analyzed on Day 5 and 9.

### Functional analysis of *in vitro* stimulated P14 CD8 T cells

The effector function and cytokine production of *in vitro* stimulated P14 cells were analyzed on Days 0, 2, 5, 7, 12, 19, 23, 26, and 30 post-initial activation. Cells were collected at these timepoints and stimulated for either 3 hours with PMA/Ionomycin plus protein transport inhibitors (eBioscience™ Cell Stimulation Cocktail, ThermoFisher), or for 5 hours with GP33 peptide (0.3 µg/ml) plus protein transport inhibitors (eBioscience™ Brefeldin A and Monesin, ThermoFisher) at 37°C. A spectral cytometry antibody targeting mouse CD107a (Brilliant Violet 421, BioLegend) was added at the beginning of each stimulation to track CD8 T cell degranulation activity. Stimulation was immediately followed by antibody staining for spectral cytometry analysis.

### Puromycin incorporation assay

For puromycin incorporation analysis (to measure protein translation capacity), P14 cells were stimulated for 5 hours with GP33 peptide (0.3 µg/ml) plus protein transport inhibitors as above. During the final 30 minutes of incubation, puromycin was added to the media at a concentration of 10 µg/ml (Millipore Sigma). During intracellular staining following stimulation, anti-puromycin antibody (Alexa Fluor 488, BioLegend) was used to label and measure puromycin uptake by spectral cytometry.

### Adoptive transfer and *in vivo* rechallenge of *in vitro* stimulated P14 CD8 T cells

For adoptive transfer experiments, antigen-experienced P14 cells were sorted on day 19 or on day 26 (following 7 days of rest) of the *in vitro* stimulation protocol to isolate the viable CD8+ T cells using the Sony MA900 Cell Sorter. Immediately following sorting, P14 cells from the three described conditions (Acute-2d, Acute-7d and T_Dysf_) were adoptively transferred into wild-type, congenically distinct (Thy1.1-Thy1.2+) C57BL/6 mice (male or female depending on P14 cell identity, age-matched at ≥ 8 weeks old) at 100-150K P14 cells/mouse from one of the three conditions via intravenous (i.v.) retro-orbital injection. One day following adoptive transfer, recipient mice were infected with LCMV-Armstrong virus (2 × 10^5^ PFU/mouse) via intraperitoneal (i.p.) injection. Mice were euthanized on day 6-7 post-infection, and lymphoid (spleens) and peripheral tissues (livers and lungs) were harvested and processed into single-cell suspensions as previously described (9). Tissue samples were then used for *ex vivo* GP33 peptide stimulation for 5 hours followed by spectral cytometry staining to assess the function and phenotype of the transferred P14 cells.

### Spectral cytometry antibody staining

Single-cell suspensions were stained with a comprehensive panel, and data were collected using a Cytek Aurora 4-laser spectral cytometer and SpectroFlow software (version 3.3). Dead cells were stained with Ghost Dye Violet 510 (Tonbo Biosciences, 1:600 dilution). Mouse surface antibodies used for staining throughout this study include Brilliant Violet 421 anti-CD107a (LAMP-1; Clone 1D4B, 1:150 dilution), Brilliant Violet 605 anti-CD62L (MEL-14, 1:400 dilution), Brilliant Violet 605 anti-CD103 (2E7, 1:400 dilution), Brilliant Violet 650 anti-Ccr7 (4B12, dilution 1:200), Brilliant Violet 711 anti-Cx3cr1 (SA011F11, 1:300 dilution), Brilliant Violet 711 Annexin V (1:300 dilution), Pacific Blue anti-Ly108 (SLAMF6; 33-AJ, 1:150 dilution), PerCP/Cyanine5.5 anti-CD8a (53–6.7, 1:300 dilution), Alexa Fluor 700 anti-CD44 (IM7, 1:400 dilution), APC/Cyanine7 anti-CD279 (PD-1; 29F.1A12, 1:200 dilution), PE/Cyanine7 anti-Tim3 (RMT3-23, 1:200 dilution), and PE anti-CD127 (Il7r; A7R34, 1:300 dilution). LCMV glycoprotein 33 (GP33) specific CD8^+^ T cells were detected by tetramerization of GP33 monomers (supplied by the NIH Tetramer Core Facility) conjugated to streptavidin-APC (eBioscience).

Cells were then fixed and permeabilized for intracellular staining using the 10X Permeabilization buffer (Invitrogen). Mouse intracellular antibodies used for staining throughout this study include APC anti-IFN-γ (XMG1.2, 1:200 dilution), Brilliant Violet 650 anti-TNF (MP6-XT22, 1:400 dilution), Brilliant Violet 785 anti-Tbet (4B10, 1:200 dilution) PE/Dazzle 594 anti-Granzyme B (QA16AO2, 1:200 dilution), PE anti-Perforin (S16009A, 1:200 dilution) Alexa Fluor 488 anti-Puromycin (2A4, 1:200) (all from Biolegend), Alexa Fluor 488 anti-TCF-7/TCF-1 (S33-966, 1:200 dilution; BD Biosciences), eFluor 660 anti-TOX (TXRX10, 1:200 dilution; eBioscience), and Brilliant Ultra Violet 737 anti-Ki-67 (SolA15, 1:150 dilution; eBioscience).

### Cytotoxicity assay of P14 CD8 T cells

For *in vitro* cytotoxicity assay, GP33-expressing CT2A glioma tumor cells were labeled with CFSE dye (1 µM, Sigma), then seeded into a 96-well flat-bottom plate at a concentration of 60K cells per well in complete RPMI medium. The tumor cells were incubated at 37 degrees Celsius and 5% CO_2_ overnight to allow monolayer formation. The following day, each well was seeded with 60K P14 cells collected from either the acutely or chronically stimulated conditions at the indicated timepoint. The co-cultured tumor cells and P14 cells were then incubated overnight (18-20hr), followed by staining for spectral cytometry. Tumor cell killing activity was determined by measuring the number of dead tumor cells per 10 viable CD8+ T cells by spectral cytometry.

### RNA sequencing

RNA was isolated from sorted P14 cell pellets using the PicoPure RNA Isolation Kit (Applied Biosystems). RNA samples were submitted for library prep and sequencing by Azenta Life Sciences. RNA-seq data was analyzed by Partek Flow software. The generated FASTQ files were used for pre-alignment QA/QC, then aligned to the mouse mm10 genome by STAR. The resultant aligned reads were quantified to the mm10 RefSeq Transcripts using Quantify to annotation model, followed by filtering to exclude features where maximum < 50.0. DESeq2 analysis was performed first by normalization of the filtered gene counts for median ratio to compare among the P14 conditions. Differentially expressed genes (DEGs) were determined as genes with fold change ≥ 2 and *p-value <0.05* between the compared groups. The DEGs were plotted into heatmaps and used for principal component analysis (PCA) compared to the transcriptional signatures of TEX subsets from acute or chronic LCMV infection in mice (28, 36) using Partek Flow Software. Gene set enrichment analysis (GSEA) for Hallmark and Gene Ontology pathways and the pan-cancer single-cell RNA-seq gene signatures for human TILs (33) was performed using Partek Flow software. The lists of significant biomarker genes for TEX subsets were calculated using the normalized RNA counts of TEX subsets (generated *in vivo* (28)) on Partek (*p-value <0.05*, fold change ≥ 2). Enrichment analyses using these transcriptional signatures of TEX subsets were performed on GSEA software (UC San Diego and Broad Institute, v4.3.3).

### ATAC sequencing

P14 cells were isolated for ATAC-sequencing using the Sony MA900 Cell Sorter, including Acute-7d or T_Dysf_ PD-1+ Tim3+ on day 19, and resting T_Dysf_ cells isolated as PD-1+ Tim3-(Rest-SP) or PD-1+ Tim3+ (Rest-DP) on day 23. ATAC was performed using Omni-ATAC, as previously described (52), with minor modifications. Briefly, nuclei were isolated from cells using ATAC lysis buffer (10 mM Tris-HCL pH 7.5, 10 mM NaCl, 3mM MgCl2, 0.1% NP-40, 0.1% Tween-20, and 0.01% Digitonin) and then transposed for 30 minutes at 37°C using Tagment DNA TDE1 Enzyme (Illumina, 20034197). Transposed DNA was purified, and libraries were generated using Nextera XT DNA Library Preparation Kit (Illumina, FC-131-2001). The concentration and size distribution of libraries were determined by Qubit (Thermo Fisher Scientific) and Agilent Tapestation, respectively. The ATAC libraries were then sequenced by Azenta Life Sciences. The data was analyzed using cutadapt 1.18, Bowtie2, Picard 2.18.17, samtools, and deepTools 3.3.0. First, the fastq files were trimmed using cutadapt, and then aligned to mouse mm10 genome assembly using Bowtie2. Next, duplicate reads were removed using Picard 2.18.17 and bigWig files were generated with deepTools. Differentially open chromatin regions (OCRs) on day 19 were defined as regions with fold change ≥2 (for Open in T_Dysf_) or ≤-2 (for Closed in T_Dysf_) and p-value <0.01 for T_Dysf_ versus Acute-7d cells and plotted as heatmaps using Partek Flow software. Principal component analysis (PCA) was performed using Partek to compare OCRs for Acute-7d, T_Dysf_ and Rest-SP or Rest-DP subsets to published chromatin landscapes for memory or exhausted CD8 T cells (23) using the top 5,000 variable peaks. To visualize chromatin accessibility transitions specific to T_Dysf_ PD-1+ Tim-3+ cells, we generated an alluvial plot. Peaks were classified as “Open” or “Closed” (Stage 2) based on comparisons to the Acute-7d condition, using thresholds of adjusted p-value < 0.01 and fold change ≥ 2 or ≤ –2. These peaks were then evaluated in resting states (Rest-DP and Rest-SP) to assign Stage 3 labels—Open, Closed, or Unchanged—using the same statistical cutoffs. The Sankey-style plot depicts the number of peaks transitioning from the Acute-7d baseline (Stage 1) through their T_Dysf_-specific state (Stage 2) to their resting fate (Stage 3), highlighting dynamic regulatory patterns across conditions. Transcription factor enrichment analysis was performed using HOMER (findMotifsGenome.pl) with the mm10 genome and the -size given option to preserve original region lengths.

### Targeted DNA methylation sequencing

DNA was isolated from FACS-purified P14 cell pellets (sorted on day 19 of *in vitro* stimulation) and used for bisulfite conversion using the EZ DNA Methylation-Direct Kit (Zymo). PCR was then performed on the bisulfite-converted DNA at differentially methylated regions for key genes related to CD8+ T cell exhaustion states (*Tcf7, Ccr7, Pdcd1*). The following primers were used for each region: *mTcf7* (Forward primer: 5’-GGTTAGTTTGAGTTTGGTTTAGAGTAGTGAG-3’, Reverse primer: 5’-CCTCTTACCTAAATTTCCCTACAAAATACC-3’), *mCcr7* (Forward primer 5’-GGAGTTTGGGATAAAAGTTTTTAATGG-3’, Reverse primer: 5’- CCAAACCCACTCTAAACCCTATATTAC-3’), *mPdcd1*(Forward primer: 5’- GGTTGAGAGAGATTGAAATTAGGGTTAG-3’, Reverse primer: 5’- CAACAAAACTAACAAACCTAAAACAACT-3’). Following PCR amplification, the amplicon size was verified by gel electrophoresis. Gel bands were excised, and amplicon DNA was purified from each region using the Zymoclean Gel DNA Recovery Kit (Zymo). The purified amplicon DNA was then used for library preparation using the Native Barcoding Kit 24 V14 (Oxford Nanopore Technologies) (9). The barcoded DNA library was loaded onto an R10 flow cell and ran on the MinION Mk1B sequencer from Oxford Nanopore. FASTQ files generated from the sequencing run were used for downstream analysis for genome alignment and determining % of CpG methylation at each amplified region using a customized NanoEM pipeline (53).

## Author contributions

Conceptualization: HEG; Methodology: HEG, AY, AAS, and AL; Investigation: HEG, AY, AAS, AL, AMY,

KP, AS, MRA and EMO; Visualization: HEG, AY, AAS, and AL; Supervision: HEG; Writing—original draft:

HEG, AY, AAS, and AL; Writing—review and editing: HEG, EMO, AY, AAS, and AL.

## Acknowledgments

We thank the NIH Tetramer Facility at Emory University in Atlanta, GA for providing the monomers used in this study for the detection of LCMV-specific GP33-tetramer+ CD8+ T cells, and Fiza Tarlochan from the Ohio State University for technical assistance. Research reported in this publication was supported by the National Institute of Allergy and Infectious Diseases of the National Institutes of Health grant R01AI170926 (HEG), the National Institute of Allergy and Infectious Diseases of the National Institutes of Health grant R01AI134035 (EMO), the National Institute of Health training program T32 AI165391 “Interdisciplinary Program in Microbe-Host Biology” (AAS), and the Ohio State University Comprehensive Cancer Center and College of Medicine funds (HEG).

