## Supplementary figures and images for "Faithful Modeling of Terminal CD8 T Cell Dysfunction and Epigenetic Stabilization *In Vitro*"

### Supplemental Fig.1

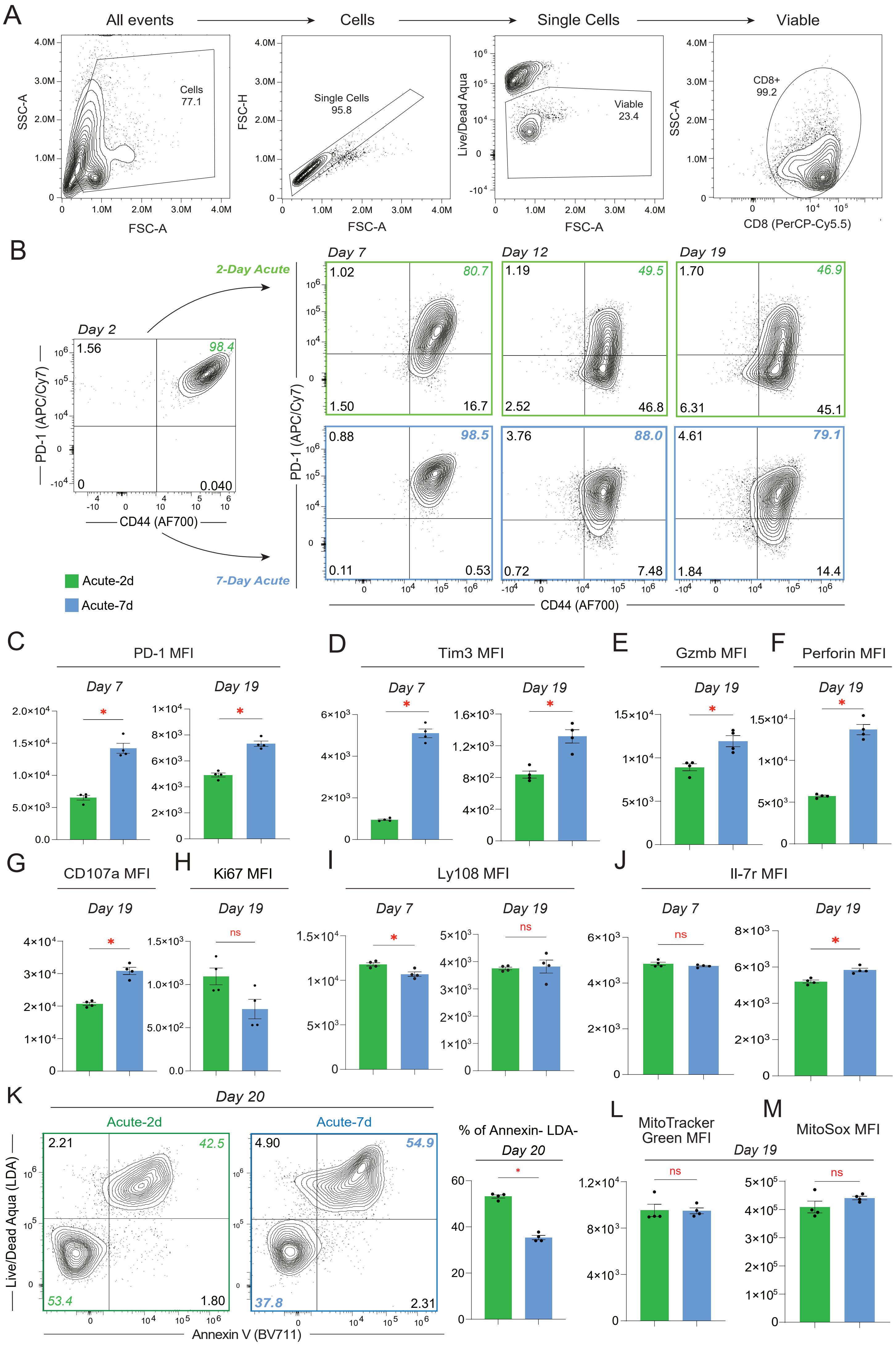


**SUPPLEMENTAL FIGURE 1**

### Supplemental Fig.2

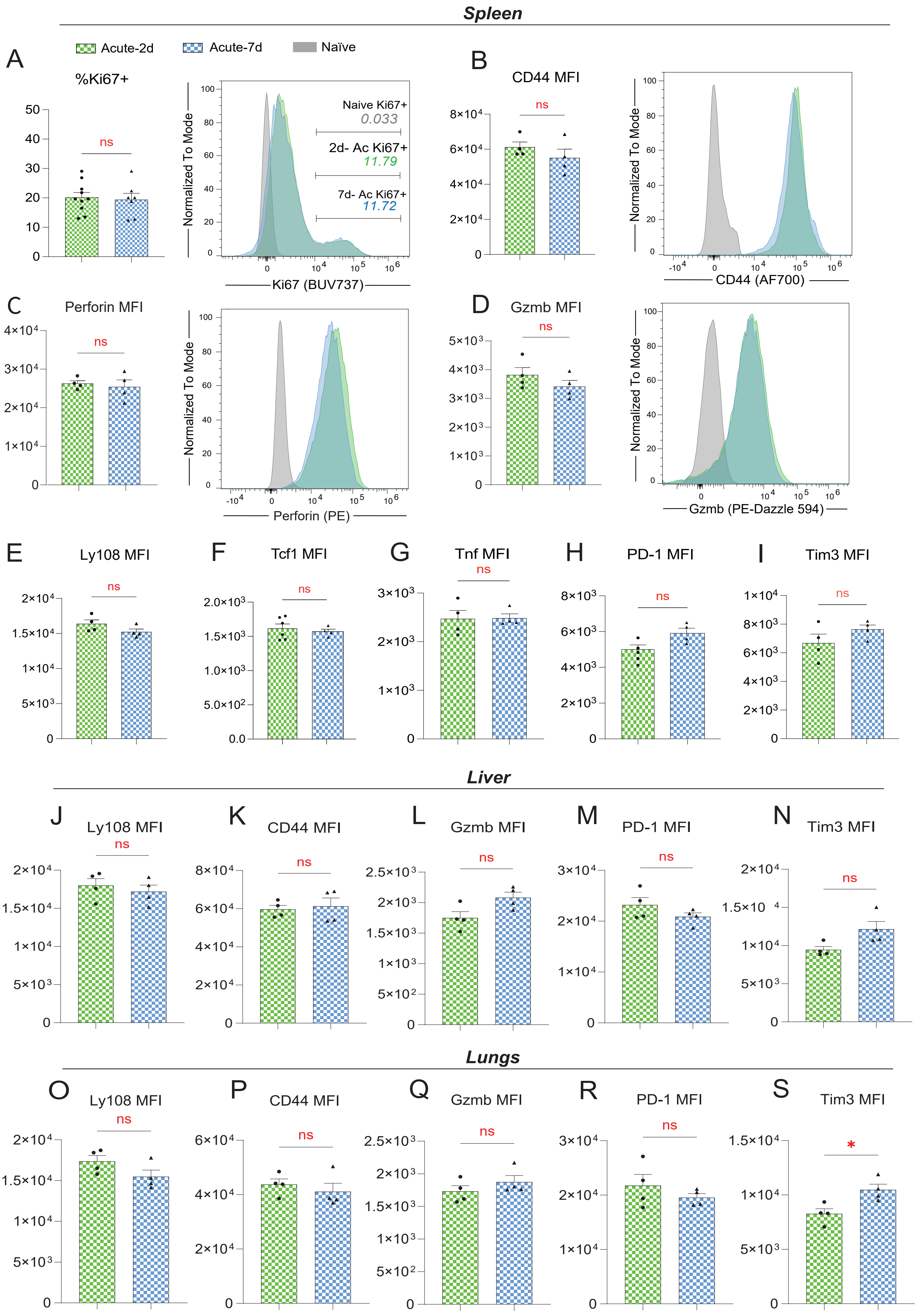


**SUPPLEMENTAL FIGURE 2**

### Supplemental Fig.3

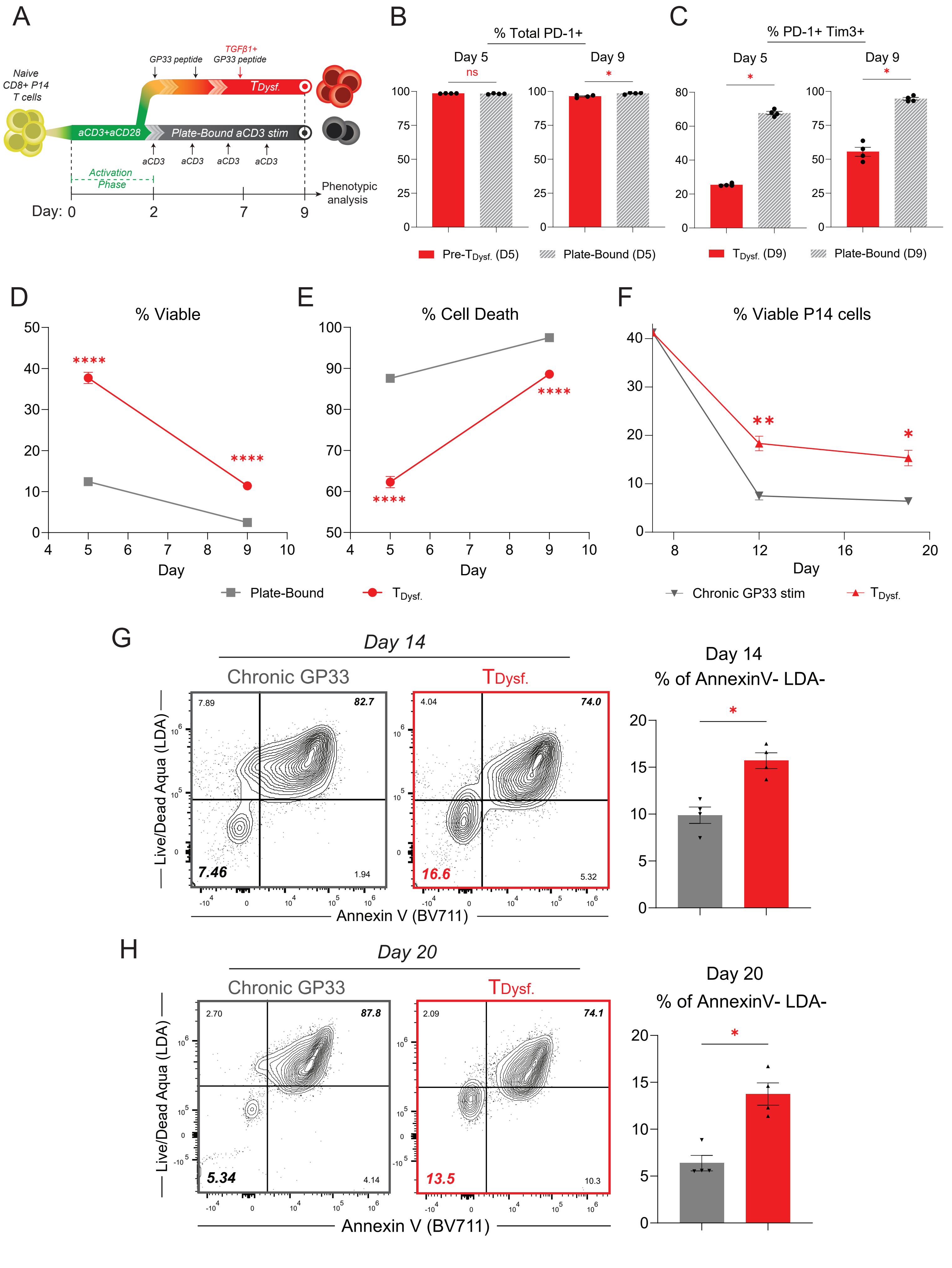


**SUPPLEMENTAL FIGURE 3**

### Supplemental Fig.4

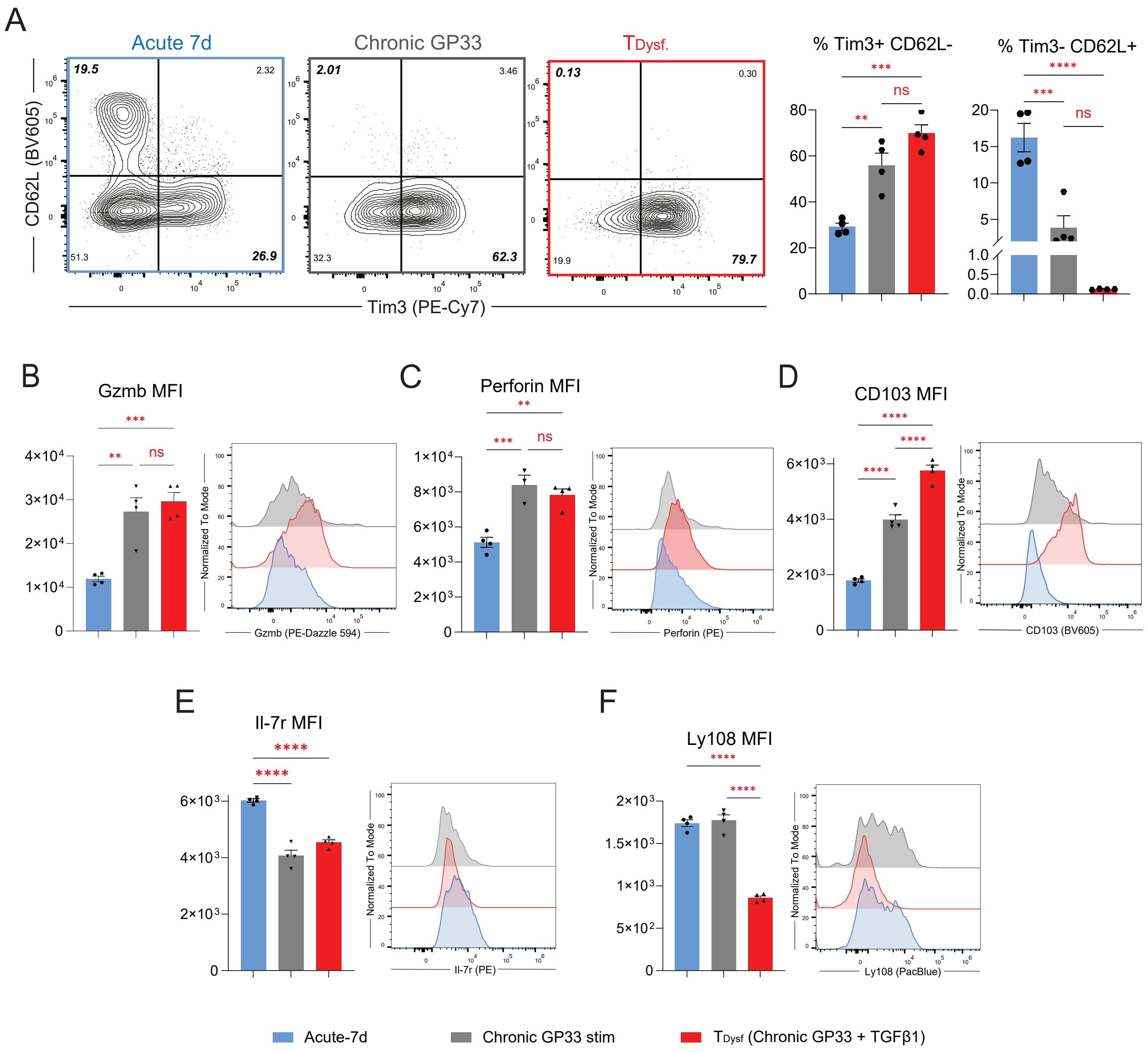


#### SUPPLEMENTAL FIGURE 4

### Supplemental Fig.5

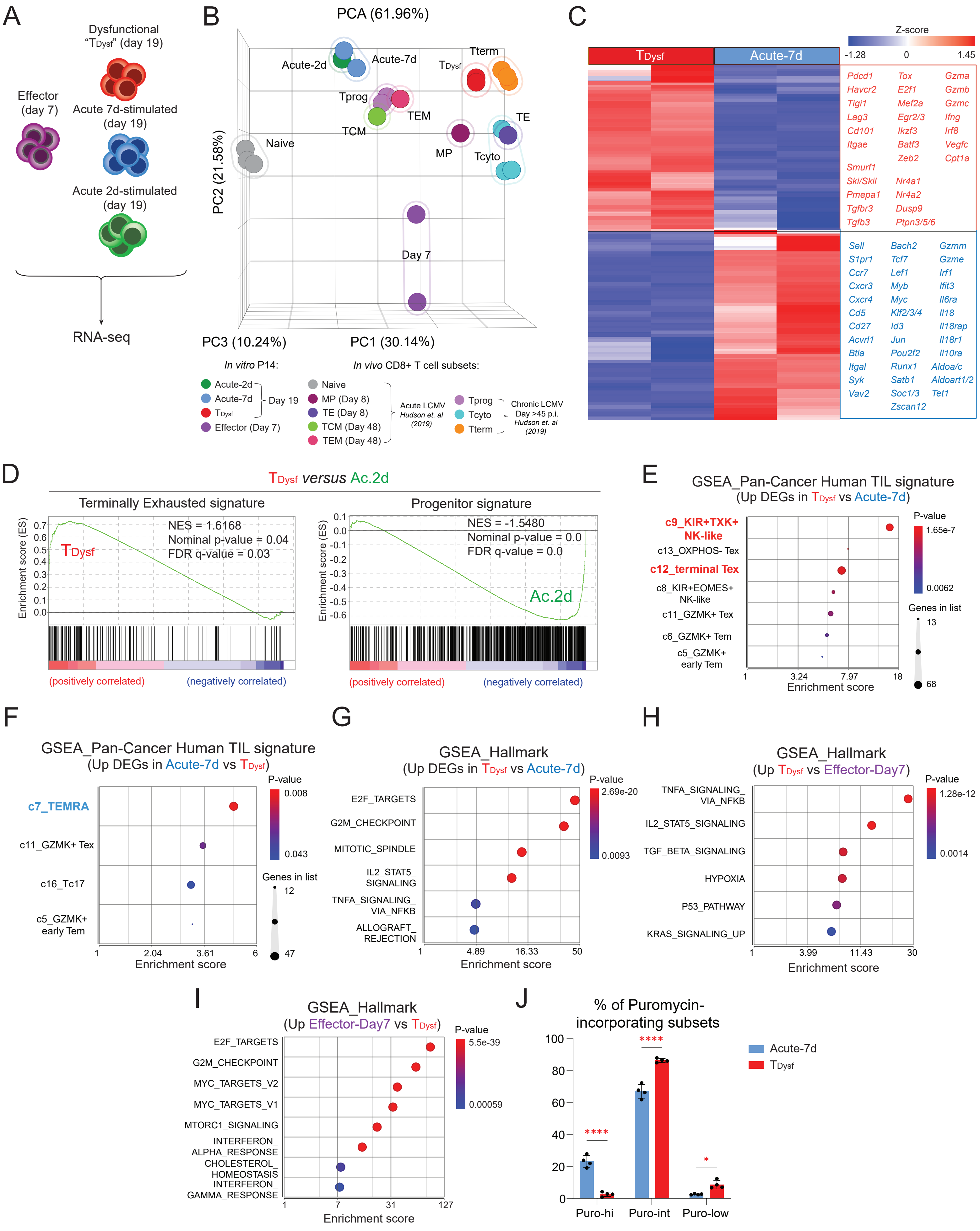


**SUPPLEMENTAL FIGURE 5**

### Supplemental Fig.6

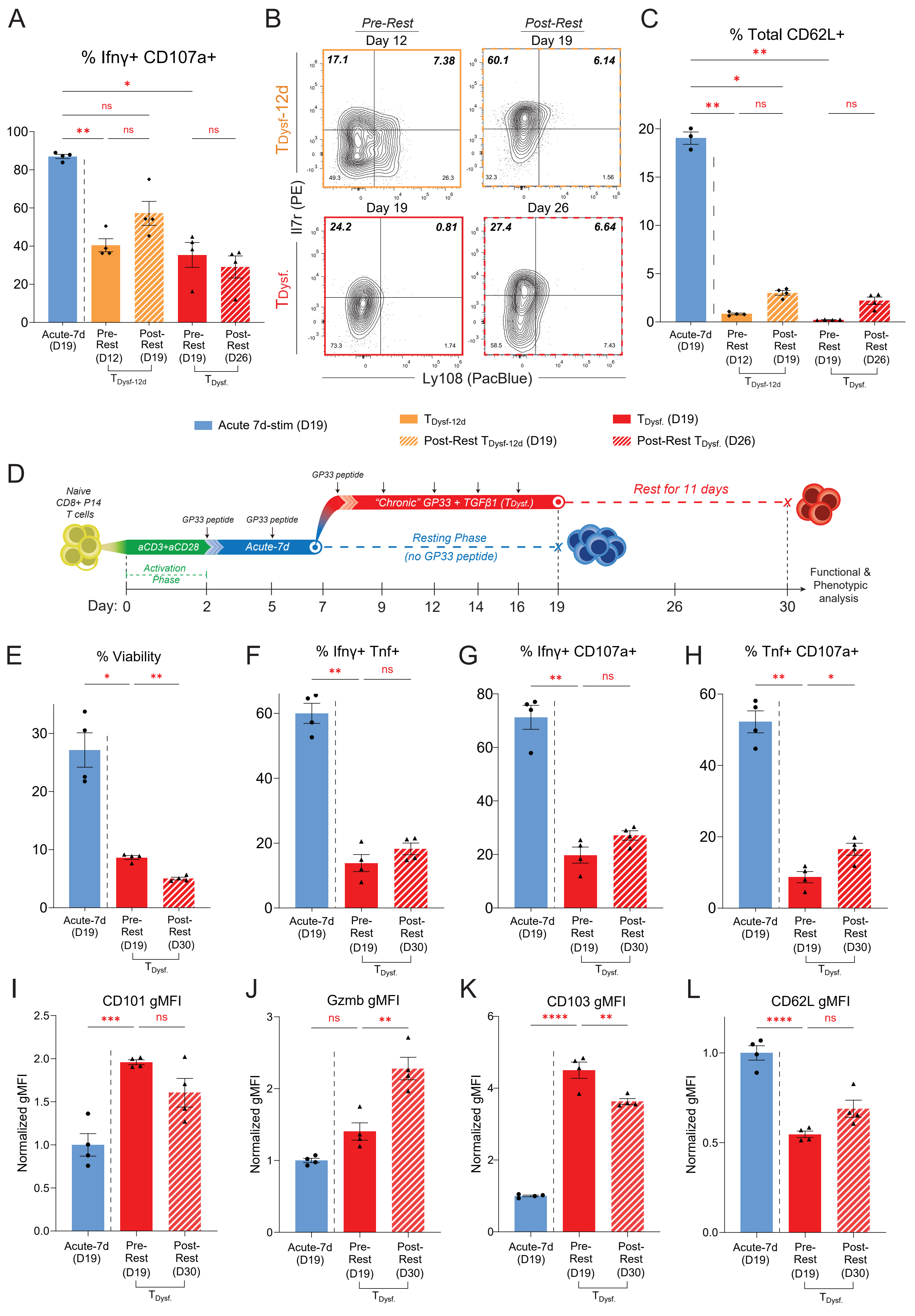


**SUPPLEMENTAL FIGURE 6**

### Supplemental Fig.7

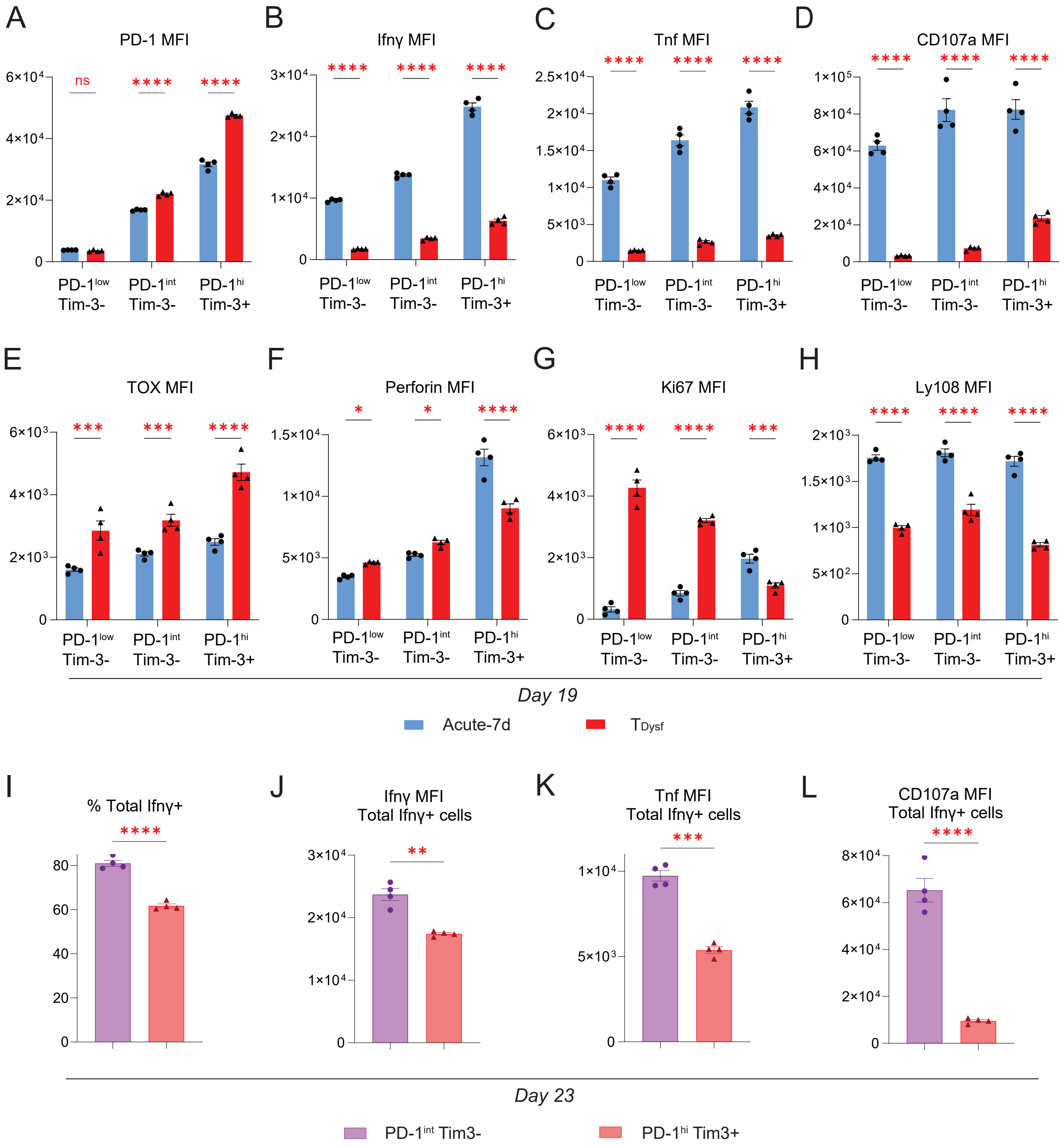


**SUPPLEMENTAL FIGURE 7**

### Supplemental Fig.8

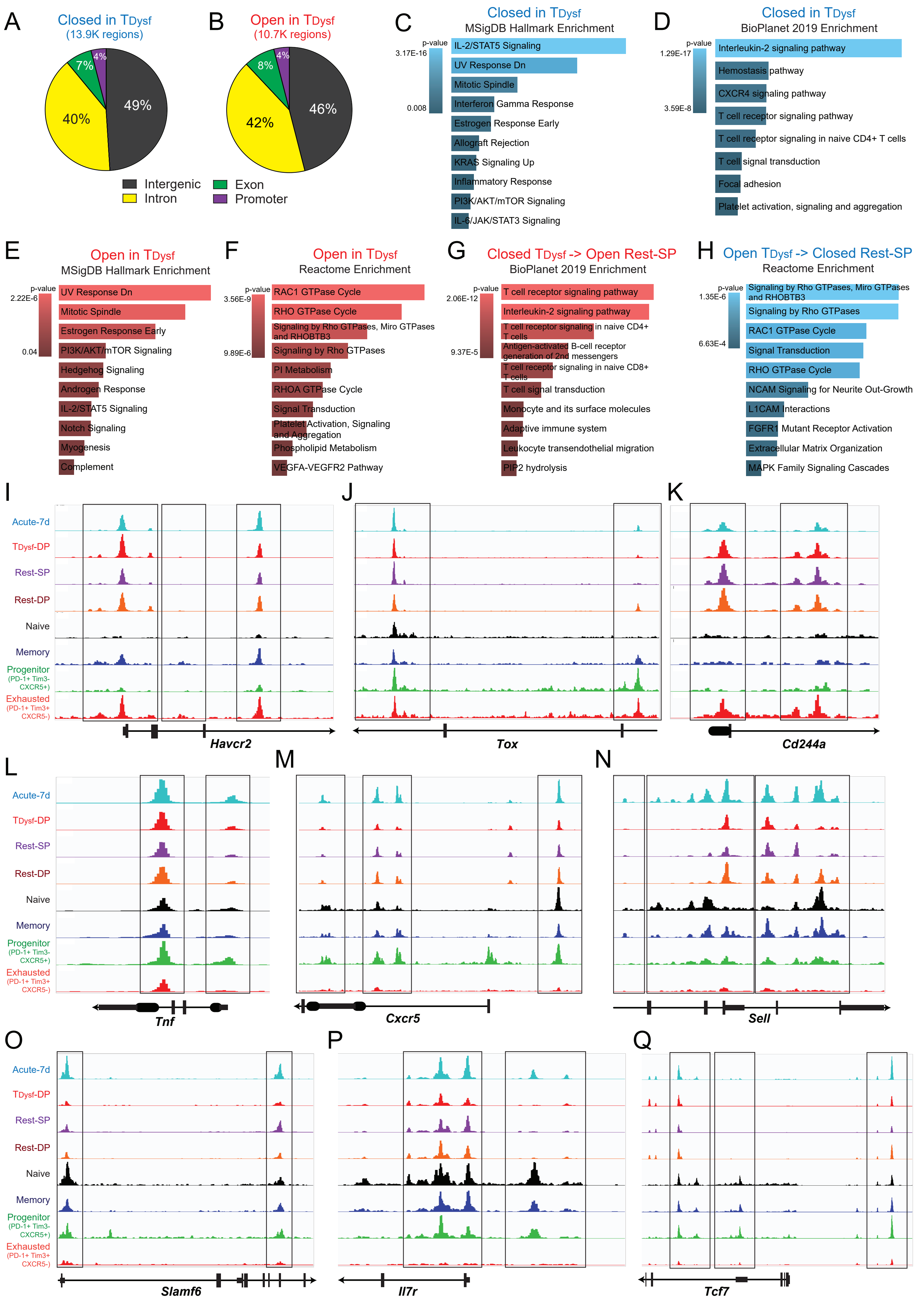


SUPPLEMENTAL FIGURE 8

### Supplemental Fig.9

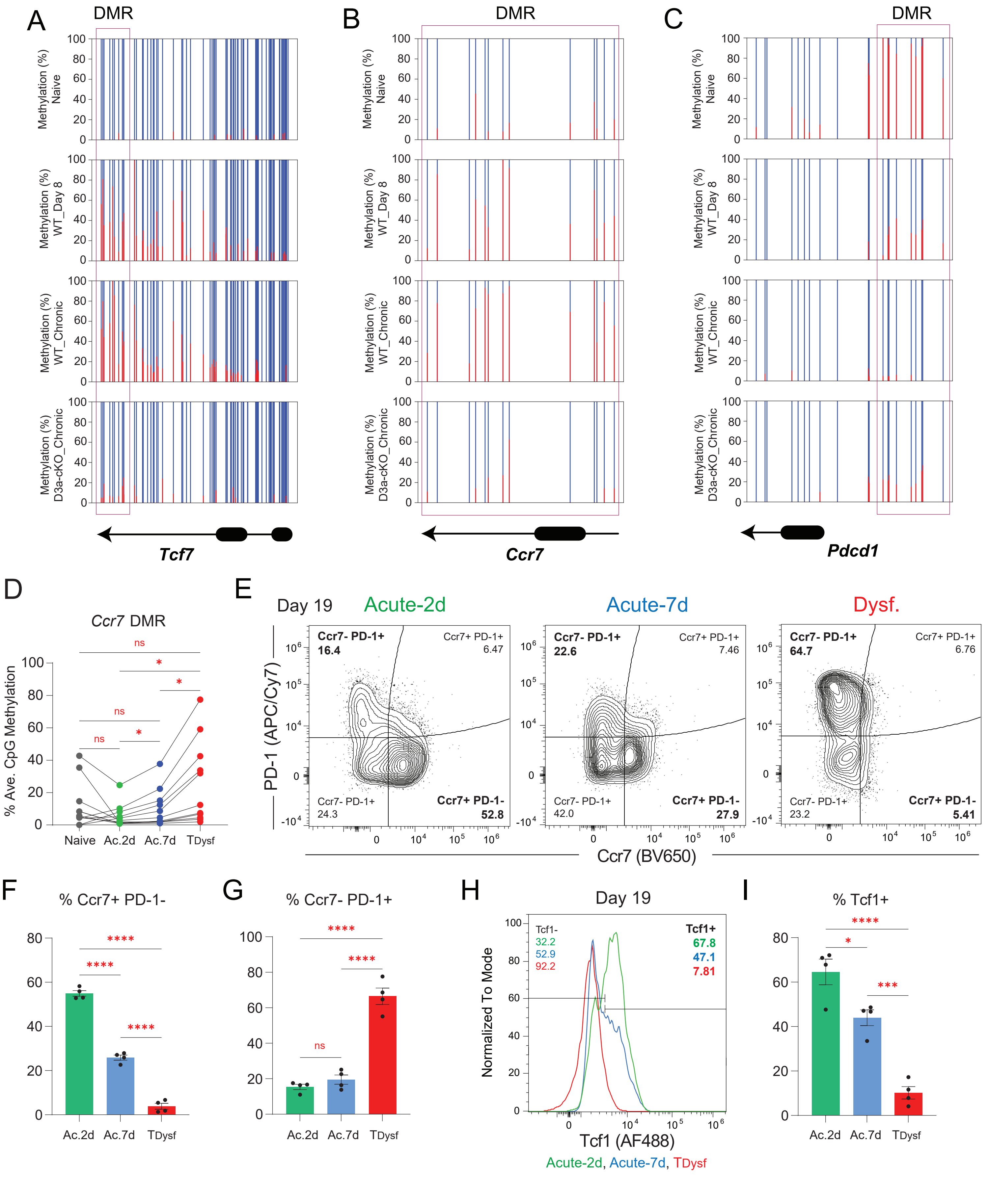


**SUPPLEMENTAL FIGURE 9**

### Supplemental Fig.10

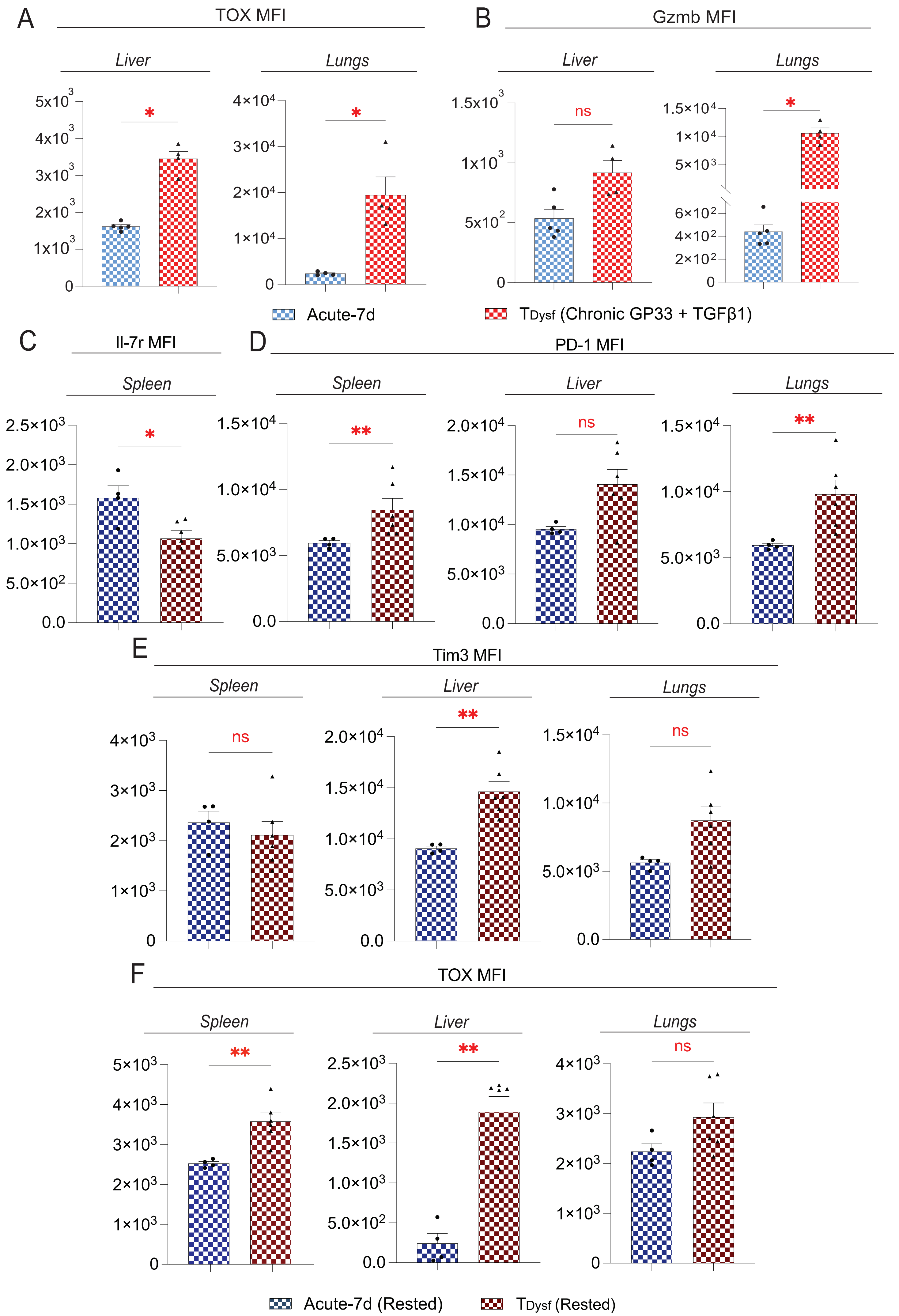


**SUPPLEMENTAL FIGURE 10**
